# *Lussac*: a fully-automated consensus method that increases the yield and quality of spike-sorting analyses

**DOI:** 10.1101/2022.02.08.479192

**Authors:** Aurélien J. G. Wyngaard, Victor Llobet, Boris Barbour

## Abstract

The rise in site counts of multi-electrodes used in extracellular recordings in the brain has driven the development of increasingly automated spike-sorting packages. However, post-processing is still largely manual and it remains difficult to determine the optimal package and parameters for a given recording. It has recently been shown that different packages produce quite disparate outputs, suggesting that a combination of analyses might be beneficial. Here, we describe the formalization of existing and new metrics of unit quality and comparison, then build upon these to automate the creation of a consensus output from multiple analyses. We validated our package against synthetic and real ground truths. Compared to individual analyses, our package increased the yield and quality of correct units (doubling the yield of Purkinje cells in our recordings) and, crucially, eliminated numerous incorrect units that were impossible to identify in a single analysis. These improvements also increase analytical objectivity and reduce manual effort.

---

Neuronal activity can be monitored by measurements of the extracellular voltage. In most analyses, multiple sources must be unmixed to recover the activity of individual neurons, in a procedure called spike sorting. The output is composed of units that, ideally, match the recorded neurons.

The advent of highly-integrated silicon probes such as Neuropixels [1] and their large volumes of data forced development to focus on increasing automation. Several efficient software suites have been developed, including Kilosort [2, 3], MountainSort [4], SpyKING CIRCUS [5], tridesclous [6] and YASS [7]. Nevertheless, spike sorting remains a difficult problem, complicated by multiple mechanisms, such as the reduction of signal-to-noise ratio with distance, waveform distortions resulting from electrode drift [1, 8] or spike collisions [9].

Although several packages brand themselves as automated, all require a manual post-processing step to remove incorrect units. This is time-intensive and its subjectivity degrades reproducibility. Furthermore, as will be shown below, package outputs sometimes contain apparently correct units that are actually the result of a merge of two or more neurons, with no way of detecting and removing them currently.

A recent survey using synthetic data has highlighted notable differences in the outputs of six spike-sorting packages [10]. Strikingly, quite large numbers of units were reported by only one package, and those units were typically incorrect, suggesting that it would be beneficial to combine the outputs of multiple packages to produce a consensus. The disparity between package outputs also highlights one of the challenges of spike sorting today: it is quite a difficult task to determine which spike-sorting algorithm performs the best on a given dataset, and what parameters to select.

We thus set out to create a procedure for combining multiple analyses that would be fully automated, minimize the subjectivity and bias of choosing a single spike sorter with a single set of parameters. The overall architecture of the resulting package *Lussac* is diagrammed in Fig. 1. Several analyses, which may differ in the spike sorter used and/or its parameters, are run on the same data. Their results, after a first, light curation, are merged to create a single output, which is curated more stringently. The main goal of this paper is to describe the construction, operation, validation and benchmarking of *Lussac*, which is based upon formalized, extended and novel metrics of unit quality and similarity. We describe the core new features in the Results section, while complete, detailed information is supplied in the Methods and appendices.

**Fig. 1.**
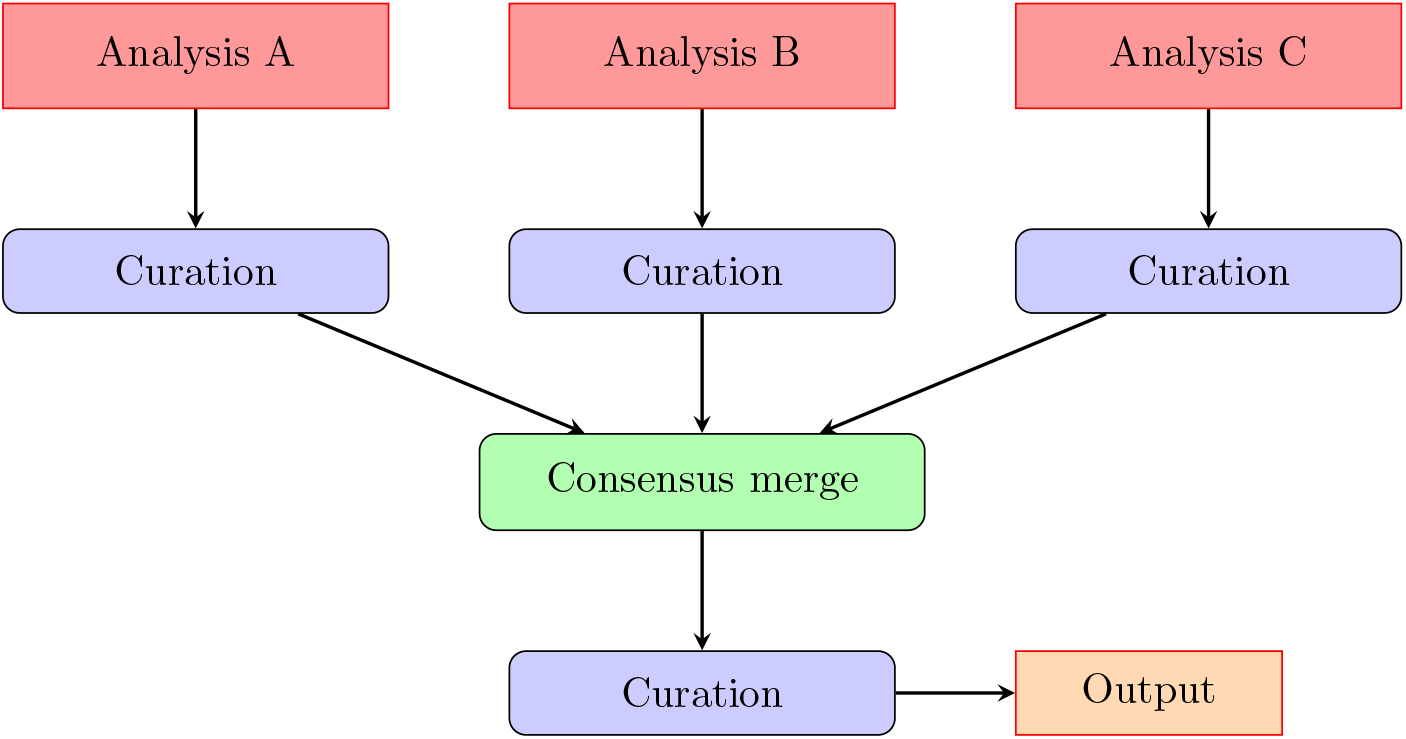
Lussac flowchart. Visual representation of the Lussac pipeline. Each analysis is run individually and curated. These intermediate outputs are then merged and curated to produce the consensus output.

In the tests shown below, we shall demonstrate that *Lussac* matches or outperforms all individual spike-sorting packages.

## 1 Results

### 1.1 Unit metrics

The creation of a consensus output requires the determination of which units from different analyses represent the same neurons, a process guided by several measures of the similarity of two units. Furthermore, quality metrics are required to decide which of two units representing the same neuron is better. Because we are the first to automate the merging of multiple analyses, the identification, formalization and implementation of these metrics must be described and justified. The first part of the Results will be devoted to describing novel intra- and inter-unit metrics, and extensions of standard metrics. A complete description of each metric is available in the Methods. When possible, we have constructed metrics with calibrated ranges, which facilitate user configuration of the package.

#### 1.1.1 Intra-unit metrics

Several metrics allow the assessment of a single unit’s intrisic quality: the Signal-to-Noise Ratio (SNR), average firing rate, contamination ratio and waveform variability. The first two metrics are well known and only detailed in the Methods. The latter two are less standardized and described below.

##### The contamination ratio

The classical criterion of spike quality is the frequency of spikes violating the (apparent) neuronal refractory period *t*_*r*_ (Fig. 2A). More informative than this number is the ratio of contaminant spikes 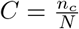 in the unit [11], where *N* = *n*_*t*_ + *n*_*c*_ the total number of detected spikes (sum of *n*_*t*_ true spikes and *n*_*c*_ contaminant spikes). This can be evaluated under the assumption of stationary firing statistics and under a model for the contaminating spikes.

**Fig. 2.**
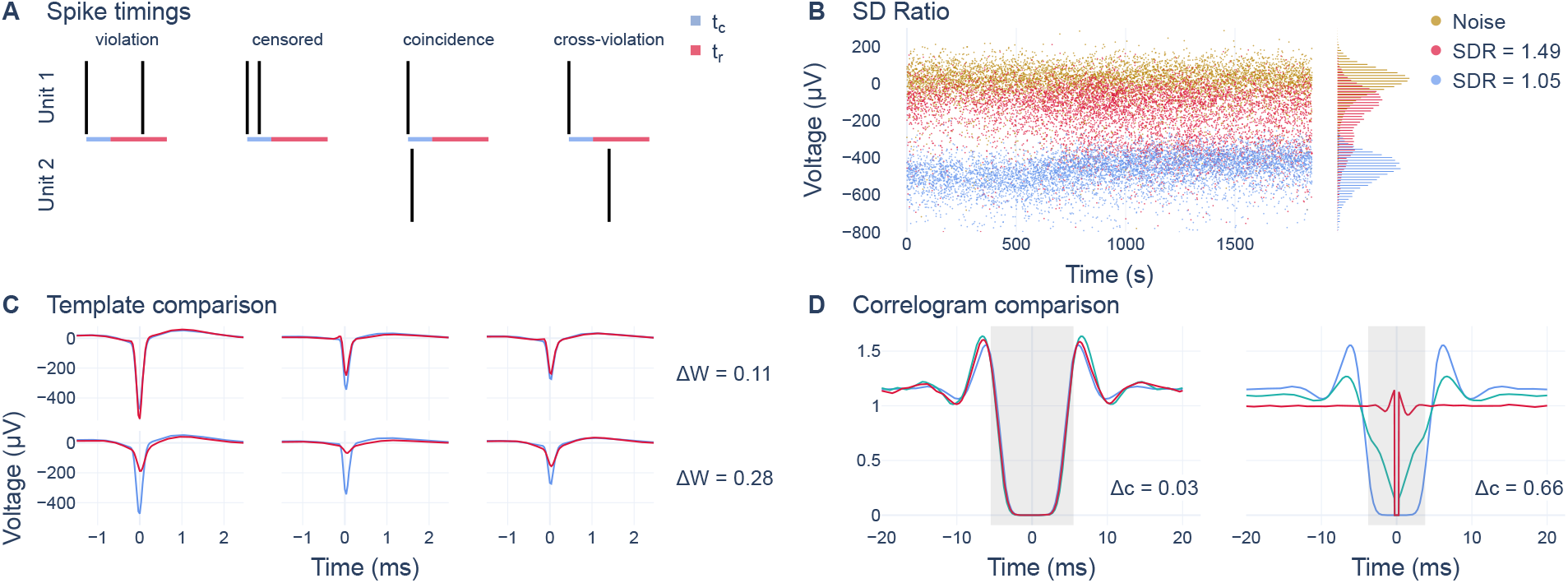
Unit metrics. (**A**) Intra-unit and inter-unit spike-timing information. From left to right: example of a refractory period violation, of censoring, of coincident spikes from two units, and of a cross-violation. *t*_*r*_ refractory period (red), which includes the *t*_*c*_ censored time (blue). (**B**) SD ratio (SDR). The spike amplitudes are shown for two units (blue, red), as well as for the baseline noise (gold), with their respective distributions on the right. The SDRs are calculated after corrections described in the main text. (**C**) Template comparison. Two pairs (rows) of units (red, blue) are shown on 3 different channels (columns). Top: pair of units from the same neuron. Bottom: pair of units from different neurons. Δ*W* waveform template difference. (**D**) Correlogram comparison. Two pairs are shown, with the auto-correlograms (blue, green) and the cross-correlogram (red). The shaded area corresponds to the comparison window. Left: same pair as panel C (top). Right: Same pair as panel C (bottom). Δ*c* correlogram difference.

Two contamination models representing extremes of a continuum can be considered. At one extreme, a single neuron is responsible for all the contaminating spikes, whereas at the other they arise from numerous other neurons (or noise). In the former case (*C*_1_), the contaminating spikes cannot violate each other’s refractory period, whereas in the latter case (*C*_∞_), they can. Given these assumptions, we can derive:

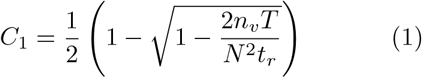

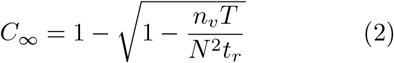

where *n*_*v*_ is the number of refractory period violations and *T* the duration of the recording. *T* and *t*_*r*_ should be understood to have been corrected for the censored period (see Methods 3.2.3). A proof for these equations is given in Appendix G.

##### Waveform variability (SD ratio)

If a unit is erroneously composed of spikes from multiple neurons, the variability of spike waveforms will, in nearly all cases, be increased compared to that of spikes from each neuron considered separately. In practice, the variation floor is set by the recording noise, which is dominated by background activity. For robustness, we constructed for each unit the simplest possible measure: the standard deviation of spike amplitudes measured at the time of the unit template’s peak, divided by the standard deviation of the noise. This is called the “SD ratio” (SDR) [12].

A value close to one is expected (implying no intrinsic spike variation), where the histogram distribution of spike amplitudes matches that of background noise (see Fig. 2B). However, several mechanisms can increase the value even for a single neuron, in particular the presence of burst spikes, which tend to be of lower amplitude, and the presence of electrode drift. We found ways to minimize the influence of both of these mechanisms on the calculation of the SD ratio (see Methods 3.2.4).

The behavior of this metric is illustrated in Fig. 2B. The blue unit representing a single neuron drifts, but the associated amplitude variations barely affect the *SDR* = 1.05. In contrast, the red unit represents a mixture of two neurons and this is readily apparent in its elevated *SDR* = 1.49.

##### Generating a unit quality score

When comparing units representing the same neuron, it is useful to have a single quality score to decide which is of better quality. Using the number of spikes and the contamination ratio, we can create a simple quality metric *Q*^′^:

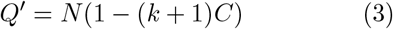

where *N* is the number of detected spikes, *C* the estimated contamination ratio, and *k* a user-defined constant balancing false negatives and false positives (see Methods 3.2.5). Throughout this paper we have used *k* = 2.5, because this was found to be close to optimal in benchmarks. It gives a weight to a false positive (incorrect spike) 2.5 times that of a false negative (missed spike).

As the SD ratio provides complementary information to the contamination (see Appendix C), the quality metric can be improved by penalizing the distance between the *SDR* and 1:

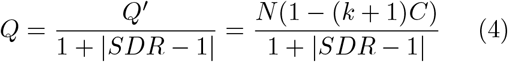

The successful application of these metrics requires that individual analyses do not bias these metrics. This problem could arise if either metric were exploited during clustering. Although this appears not currently to be the case with the packages tested here, an analogous bias can be introduced indirectly, probably as a result of detection problems caused by spike collisions. For example, as shown in Appendix C, MountainSort produces units whose apparent contamination is lower than the real value. We speculate that because the package does not perform template matching to redetect spikes after clustering, some spikes occuring during the refractory period are lost during clustering and not recovered, leading to an under-estimation of the contamination.

##### Undetectable merges

Although many incorrect units can be removed using the above metrics, seemingly high-quality units can result from merges of two neurons or other mechanisms. Some examples from our synthetic dataset used below are shown in Appendix D. Manual post-processing on the basis of a single analysis struggle to detect such incorrect units, which is a pernicious failure of spike sorters that can lead researchers to mistake a merge for a high-quality unit.

However with multiple analyses, additional information is potentially available, as only one analysis needs to correctly split the neurons into different units for the error to become detectable. Exploiting this information is a key innovation of the *Lussac* package, and allows removal of most of these seemingly high-quality erroneous units. This makes use of inter-unit metrics.

#### 1.1.2 Inter-unit metrics

The automatic exploitation of information from multiple analyses requires the identification of the units from different analyses that are likely to represent the same neurons. We developed and implemented specific pairwise similarity metrics to guide these decisions: the spike coincidence (see Methods 3.2.6), template comparison, correlogram comparison and cross-contamination. As will be seen below, the units of multiple analyses can be represented as the nodes of a graph, while the pairwise metrics listed above, attached to the edges between each pair of nodes, characterize their similarity. Analysis and simplification of this graph will then generate the consensus output.

##### Template comparison

A key condition for two units to represent the same neuron is that their template (average) waveforms *W* must be similar (see Fig. 2C). The difference Δ_*W*_ between two templates *A* and *B* is computed as the ratio of of the absolute value of the differences to the total signal:

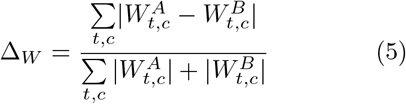

where the difference is computed along time *t* and across multiple channels *c*. This metric ranges between 0 (identical templates) and 1.

##### Correlogram comparison

Auto- and cross-correlograms offer powerful diagnostics for identifying units belonging to the same neuron and are usually used by researchers during manual post-processing. We thus set out to automate this procedure. A noteworthy property is that the expected shape of a correlogram is unchanged if spikes are removed *randomly* (a proof is given in Appendix H): two daughter units generated by a random split will have auto- and cross-correlograms similar (but noisier) to the auto-correlogram of the parent unit.

Correlograms between all pairs of units are computed, filtered to reduce binning noise and then normalized (see Fig. 2D). The differences of the auto-correlograms with the cross-correlogram were calculated as the average of the absolute differences. The correlogram difference Δ_*c*_ was computed as a weighted sum of both differences:

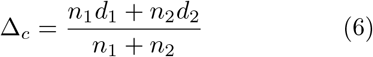

where *d*_1_, *d*_2_ are the differences between the cross-correlogram and each auto-correlogram, and *n*_1_, *n*_2_ the numbers of spikes in each unit. These comparisons were restricted to a region around the central dip caused by the apparent refractory period (see Methods 3.2.8). This maximizes the sensitivity of the comparison, as this central region displays the highest variability between neurons.

##### Cross-contamination

Relevant information to identifying erroneous seemingly high-quality units is the rate at which spikes from one unit violate the refractory period of another unit (from a different analysis) known from spike coincidences to represent, at least in part, the same neuron. Such “cross-violations” (Fig. 2A) can only occur at a high rate if at least one of the units contains spikes from multiple neurons.

We constructed a metric quantifying cross-violations and term this the “cross-contamination”. This metric is directional: the cross-contamination *CC*_*B*→*A*_ represents the fraction of spikes in unit *B* that do not correspond to the neuron represented by unit *A* (this fraction may arise from another neuron, or many neurons):

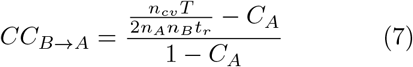

where *n*_*A*_ and *n*_*B*_ are the number of spikes in each unit, *C*_*A*_ the (estimated) contamination of unit *A, T* the duration of the recording, *t*_*r*_ the refractory period, and *n*_*cv*_ the number of cross-violations. In this dimensionless metric, 0 corresponds to the situation in which the units represent the same neuron, whereas 1 corresponds to the case of units representing two different uncorrelated neurons.

In this model, *n*_*cv*_ can be very well approximated with a binomial distribution, which we use to create a statistical test for more robust comparisons, mainly to limit absurd results due to low number of spikes (see Methods 3.2.9).

### 1.2 The *Lussac* pipeline

The first step of applying the consensus method is to run multiple analyses. These will typically involve diverse packages and/or different parameters. These analyses can be ran through *Lussac*, whose flexible pipeline (Fig. 1) applies parameters read from a unified configuration file and then works automatically and sequentially through different modules to curate and merge the analyses. All *Lussac* modules are described in depth in the Methods: unit alignment, unit annotation, removal of duplicate spikes, removal of low-quality units, removal of redundant units, intra- and inter-analysis merges. We shall outline here the intra- and inter-analysis merges, as these are the core and most innovative parts of *Lussac* in the creation of a consensus of multiple analyses.

#### 1.2.1 Intra-analysis unit merge

Modern spike-sorting packages tend to generate too many rather than too few clusters. This is a valid policy, as it is much more difficult to detect and split incorrectly merged units than it is to merge units of an incorrectly split parent. A significant portion of manual post-processing involves the identification and merging of incorrectly split units, which *Lussac* automates. However, as *Lussac* can exploit additional information at the stage of combining multiple analyses, this intra-analysis merge deliberately only combines clear-cut cases.

The merging process relies on finding pairs of units with low template and correlogram differences, and for which a merge produces a unit with a higher quality score than those of both original units (see Methods 3.3.6).

#### 1.2.2 Inter-analysis unit merge

The production of a consensus between multiple analyses is quite complex (Fig. 3). The first step is to construct a graph of all units from all analyses, linking units potentially representing the same neurons. The graph is then progressively refined to isolate groups of units representing the same neuron for which, finally, a consensus output for each neuron is generated.

**Fig. 3.**
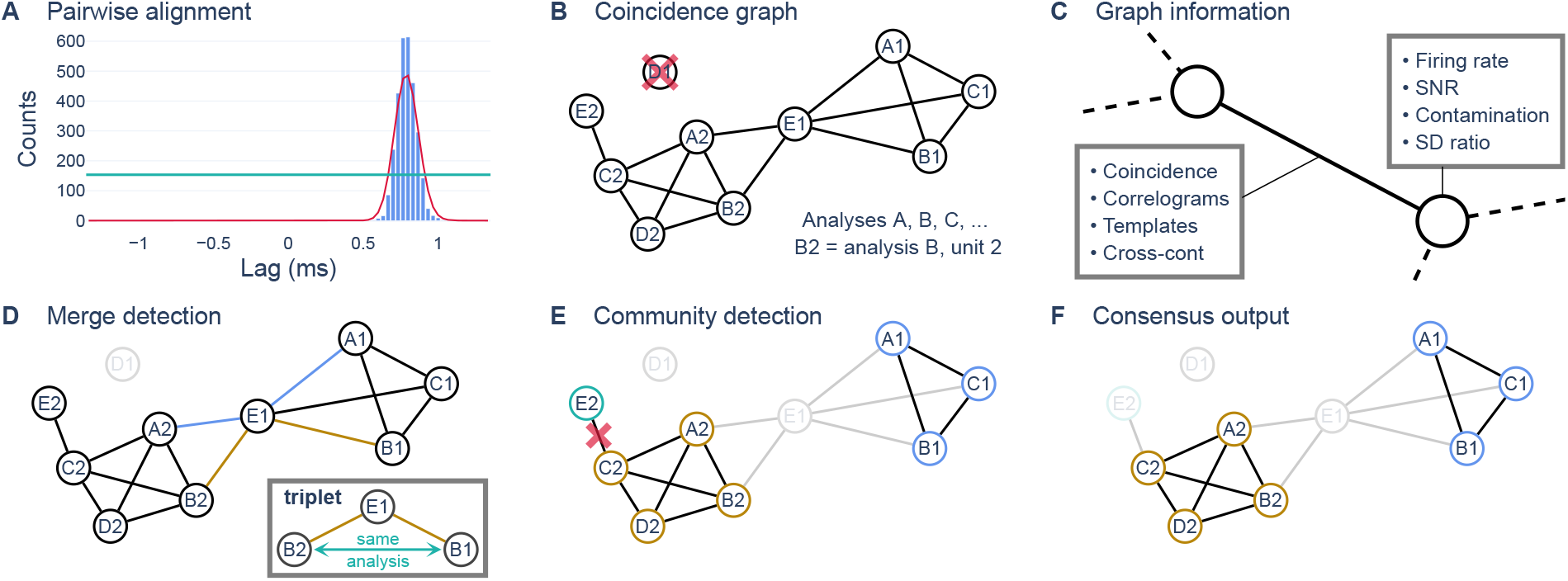
Graph analysis method of inter-analysis unit merge. (**A**) To identify the correct mutual alignment for every pair of units, the cross-correlogram (blue) is computed then filtered (red). If the peak is higher than the threshold (green horizontal line), then the correct shift is at the time of the peak. (**B**) Units are represented as the nodes of a graph and connected depending on their similarity quantified by the coincidence of their spike trains. Singletons (e.g. D1) are removed. (**C**) Representation of available nodal and edge information. (**D**) “Triplets” of two units from the same analysis indirectly connected by a node from another analysis (e.g. blue, gold edges; inset) are detected, and merged units are removed. (**E**) Communities are detected (each color is a different community) and edges between communities are removed. (**F**) The final, refined graph is used to produce the consensus output.

##### Pairwise aligment

A crucial technical preliminary to any pairwise comparison between units is to ensure that their waveforms are mutually aligned. This was done by computing and filtering the cross-correlogram (see Methods; Fig. 3A). A thin, high peak is expected if units are very similar, and the timing of the peak corresponds to the time shift between them, yielding the correct alignment.

##### Creating the similarity graph

In the graph, each unit (represented as a node) is connected to another unit by an edge if their similarity of coincident spikes is at least 30 % (see Methods; Fig. 3B). This yields a graph linking units potentially representing the same neuron. By default, unconnected nodes are removed because they cannot contribute to a consensus and are usually garbage units [10] (although in some cases, keeping them is beneficial).

The nodes and edges are populated with the intra-unit and inter-unit metrics described above and in the Methods (Fig. 3C). Thus, every node has 4 quality metrics, every edge has 4 similarity metrics and every pairwise comparison of units can exploit information in 12 dimensions.

##### Removal of merged units

The first major step of refining the graph addresses a problem that is almost insoluble within a single analysis: the identification and removal of merged units. In the graph, such units form a subgroup of nodes like E1 (from analysis E) in Fig. 3D, connected to nodes B1 and B2 (from another analysis, B), forming an incorrect link between two neurons. Evaluation of all such triplets to identify and remove those that represent merged units is a crucial step.

Two main scenarios can explain triplets: either the central node (E1 in our example) is the result of a merge of multiple neurons, or the terminal nodes result from an erronerous split. To distinguish the two cases, we developed the cross-contamination metric (see section 1.1.2). A low cross-contamination between the terminal nodes is expected if they arise from a split, whereas a high value is expected if the central node is an erronerous merge. Thus, if the cross-contamination is significantly higher than a certain threshold, the central node is removed from the graph.

##### Removal of bad connections

Each edge is checked individually, using a function of all of the inter-unit metrics. Edges failing to pass the composite similarity thresholds are removed, and any node that becomes disconnected in this process is also removed.

Furthermore, since the removal of such edges implies they were errorneous, the quality of both nodes is re-evaluated using more stringent thresholds for the contamination and SD ratio metrics. If these criteria are not satisfied, the node is removed.

##### Community detection

So far refinement of the graph has made use of information only from single nodes or edges. However, additional information is available if the whole graph is considered. In particular, if many analyses correctly find a neuron, their nodes are likely to have locally dense connections in the graph. This property is used to identify groupings of units representing the same neuron. In graph analysis, such groupings are called communities (Fig. 3E) and many algorithms exist for identifying them.

We use the Louvain community detection algorithm, without any weighting of the edges (i.e. only looking at connectivity). All edges that link nodes from two different communities are removed. Any resulting communities of size 1 or 2 are also removed. If *Lussac* was run with less than 4 analyses, this entire step is skipped, because not enough information is available.

##### Creating the consensus output

At this stage, the graph has been refined and each community should correspond to a neuron. The aim now is, for each community, to output the best representation of its putative neuron. For each node in the community, the quality metric (Eq. 4) is computed, and the unit with the highest score is identified.

The algorithm then creates a “consensus unit”, containing the spikes that were detected by at least *n* analyses (by default, 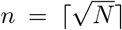, where *N* is the number of nodes). If the quality score of the consensus node (Eq. 3) exceeds the quality of the best node (∼ 81 % of cases in the synthetic benchmarks used below), then it is used for the consensus output.

### 1.3 Benchmarks

To test the performance of *Lussac* and other algorithms, we developed several validation methods using a synthetic dataset, a hybrid dataset created by injecting waveforms into real data and finally by exploiting a unique feature of the cerebellum to deduce a ground truth in real cerebellar cortical recordings.

#### 1.3.1 Synthetic neo-cortical datasets

Synthetic datasets have the major advantage of providing an exhaustive ground truth. We used the MEArec package [13], which simulates the extracellular voltages resulting from model neo-cortical layer 5 pyramidal cells and inteneurons. To make this dataset as realistic as possible, we simulated 3,000 neurons, most of which are unrecoverable (see Fig. 4A-B). This choice was made to mimic the distribution of neuronal SNRs and the biological noise, which arises in large part from the neurons that are too distant to detect. For a detailed description of the dataset, see Methods 3.4.1. Two datasets were generated: a stationary one with no electrode drift, and the other with both fast and slow drift (see Fig. 4C).

**Fig. 4.**
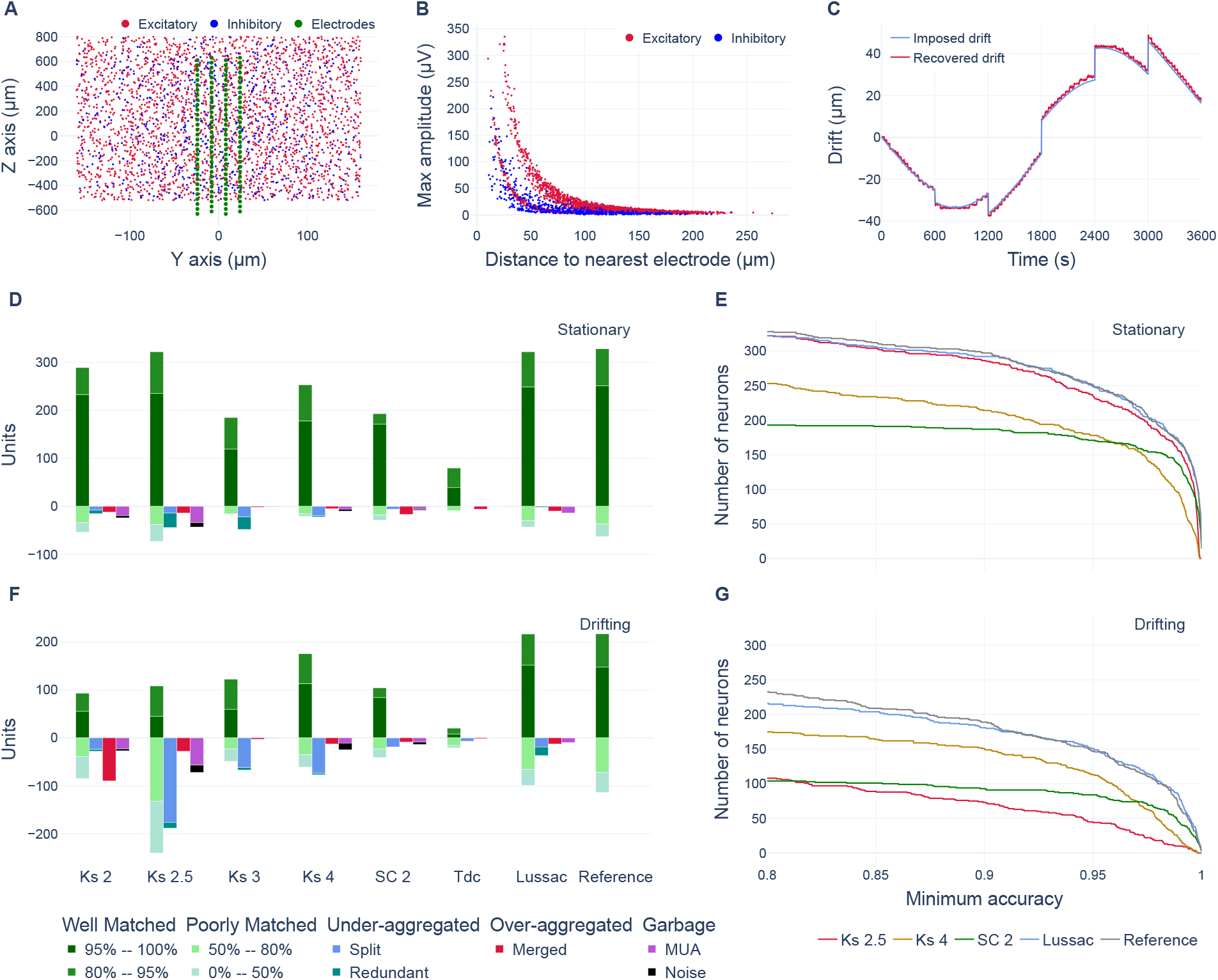
Synthetic neo-cortical benchmarks. (**A**) Spatial distribution of synthetic neurons relative to the electrodes (truncated Neuropixels 1 layout) in the YZ plane. (**B**) Distribution of template maximum amplitude vs. distance of neuron from the nearest electrode. (**C**) Electrode drift used in the drifting recording (Z axis). Drift is discretized by MEArec, as observed from its estimation (red) by the SpikeInterface algorithm [8, 10]. (**D**,**F**) Bar plots showing the classification of every output unit for each analysis for the stationary (D) and drifting (F) datasets. Units classified as good are counted upwards (above *y* = 0), whereas units classified as bad are plotted downwards. Matched units are classified based on their accuracy. MUA stands for Multi-Unit Activity. (**E**,**G**) For each analysis, the number of neurons recovered with an accuracy greater than or equal to the value on the *x*-axis is plotted for the stationary (E) and drifting (G) datasets. Unit accuracy is defined in Methods 3.4.3. The reference is constructed by taking the best unit for each neuron from all individual analyses.

We ran multiple spike-sorting packages with default parameters: Kilosort 2/2.5/3/4, SpyKING CIRCUS 1/2 and Tridesclous. For packages that had not implemented drift correction, we used SpikeInterface’s method for drift correction. For some packages where notable gains were observed when parameters were adjusted, additional analyses were also included. For each analysis, automated post-processing removed obviously garbage units (see Methods 3.4.2). No further processing was applied. We then automatically classified units using the ground truth data (see Methods 3.4.3). The joint objectives were to maximize the number of well-matched units recovered, maximize their accuracy *and* minimize the number of errorneous units. The results are shown in Fig. 4. A selection of the 11 analyses are shown here; all are presented in Appendix A.

The *Lussac* run combined all the 11 individual analyses. We also ran a reduced version combining only 4 pre-defined analyses (those we would have chosen had we been limited to 4 analyses), which yielded similar results (see Appendix A).

For the stationary dataset, we found that Kilosort 2.0 and 2.5 were the individual analyses that recovered the most neurons after parameter tuning, although at the cost of significant numbers of problematic units in the output (Fig. 4D). For the more difficult drifting dataset however, Kilosort 4.0 took the lead, although it too reported several problematic units (Fig. 4F). SpyKING CIRCUS 2 performed well with high-accuracy units (≥ 95 %) and reported few problematic units both in the stationary and drifting cases. However, this performance came at the cost of recovering fewer units with moderate accuracy < 95 % (Fig. 4E,G). In general, it can be seen that all individual packages are unable to recover large numbers of units without including significant numbers of erroneous units. *Lussac* managed largely to avoid this trade-off and outperformed every individual analysis in the number of neurons recovered with a very high accuracy, while outputting fewer problematic units, both on the stationary and drifting datasets. We can see that on the more challenging, drifting dataset, there is a significant gain when using *Lussac*, with a benefit of at least 33 % more neurons recovered with a minimum accuracy of 95 %, associated with a 40 % reduction in the amount of merged, MUA and noise units compared to Kilosort 4 with optimized parameters.

#### 1.3.2 Real cerebellar cortical datasets

Multielectrode data recorded in the cerebellar cortex is typically challenging for spike sorting because of the wide variety of firing rates and signal frequencies in the spikes. Simple spikes are narrow and emitted at high rates, whereas complex spikes are longer-lasting events with low-frequency components and occur at much lower rates. This makes the cerebellar cortex an interesting test case for a consensus method, where different analyses can be optimized for different spike waveforms.

The data presented here were obtained from chronic recordings in experiments studying changes of cerebellar cortical activity during eyeblink conditioning. The data and recording methods are described in Methods 3.4.4.

We assessed the performance of the different packages in two different ways. The first exploits the unique characteristics of the Purkinje cell, in which two types of spike must be observed, as a form of ground truth. The second uses spike injection to evaluate the recovery of spikes from different cell types.

##### Recovery of Purkinje cells

Simple and complex spikes of Purkinje cells have different waveforms and are therefore usually sorted into distinct units (see Appendix E). One can easily link the simple and complex spikes of the same Purkinje cell by examining their templates (which are very similar after high-pass filtering) and cross-correlogram (in which the complex spike causes a profound pause of simple spike firing, see Methods 3.4.5). This combination of features (simple spike, complex spike, waveform similarity, pause) is very unlikely to occur except when a Purkinje cell has been faithfully recorded. Thus, the recovery of a Purkinje cell constitutes a reliable ground truth.

Fig. 5B shows the recovery of simple/complex spike pairs for different analyses. *Lussac* yielded more than 80 % Purkinje cells than the best individual analysis and the current gold standard.

**Fig. 5.**
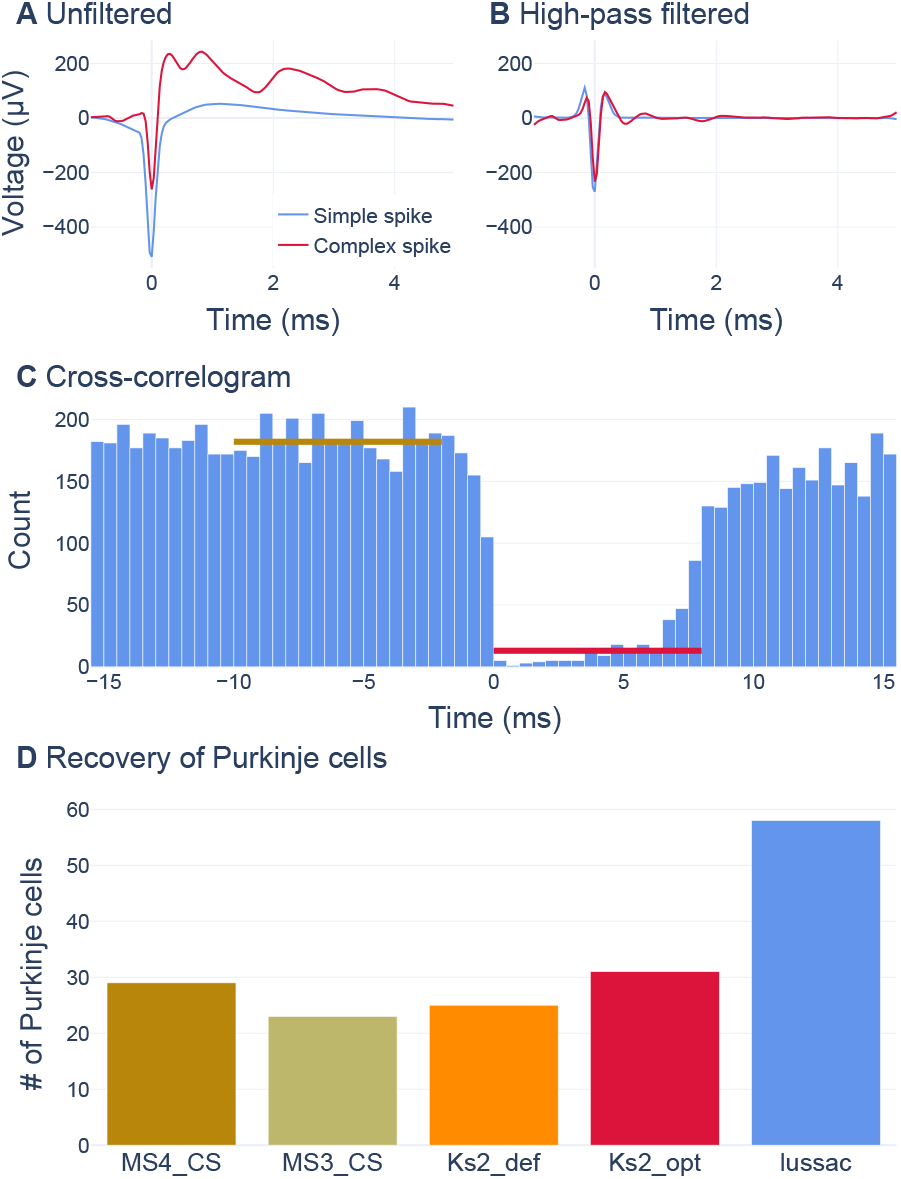
Recovery of Purkinje cells in real recordings. (**A-B**) Example of a complex and simple spike average waveform from the same Purkinje cell, unfiltered (A) and high-pass filtered at 1500 Hz (B). (**C**) Example of a cross-correlogram between the complex and simple spikes shown in (A-B), with the characteristic “pause” in simple spike firing after the complex spike. Gold line: baseline ; Red line: median value between 0 and 8 ms. (**D**) Quantification of the recovery of Purkinje cells with both simple and complex spikes for several analyses. Purkinje cells were defined as cross-correlograms with a red line lower than 40 % of the baseline.

The recovery of complex spikes within cerebellar activity is a situation where a consensus method is particularly beneficial. Indeed, many spike-sorting packages are poorly optimized for the long-lasting waveforms of complex spikes and such optimization might degrade performance for other types of spike.

##### Spike injection

To test the various packages on different cerebellar cell types, we created hybrid datasets as follows. A set of templates was created from well-detected units (without using *Lussac*, to avoid a bias in favour of our package) and then injected into recordings from different mice in scaled-down versions, in the correct cerebellar cortical layers (see Methods). These hybrid datasets were then reanalyzed to quantify the recovery of the injected units. The analyses ran were Kilosort 3/4 and SpyKING CIRCUS 2 (default parameters), Kilosort 2 (CS optimized and SS optimized), Kilosort 2.5 (optimized), and Mountainsort 3/4 (CS optimized).

The limitations of such a technique, which should be borne in mind when evaluating the results, are that the templates are stationary, there is no intrisinc waveform variability, and the complex geometry of underlying cells may not be respected (such as the parallel planar dendritic trees of Purkinje cells). This is likely to make the recovery of injected units easier than real ones. For simplicity, we also injected Purkinje cell simple and complex spikes separately.

In terms of the accurate recovery of neurons, the results in Fig. 6A-D show that *Lussac* matched or outperformed every individual analysis, regardless of the cellular type. The gap was more pronounced for complex spikes. For clarity, not all analyses are shown in the figure, but the complete set is shown in Appendix B.

**Fig. 6.**
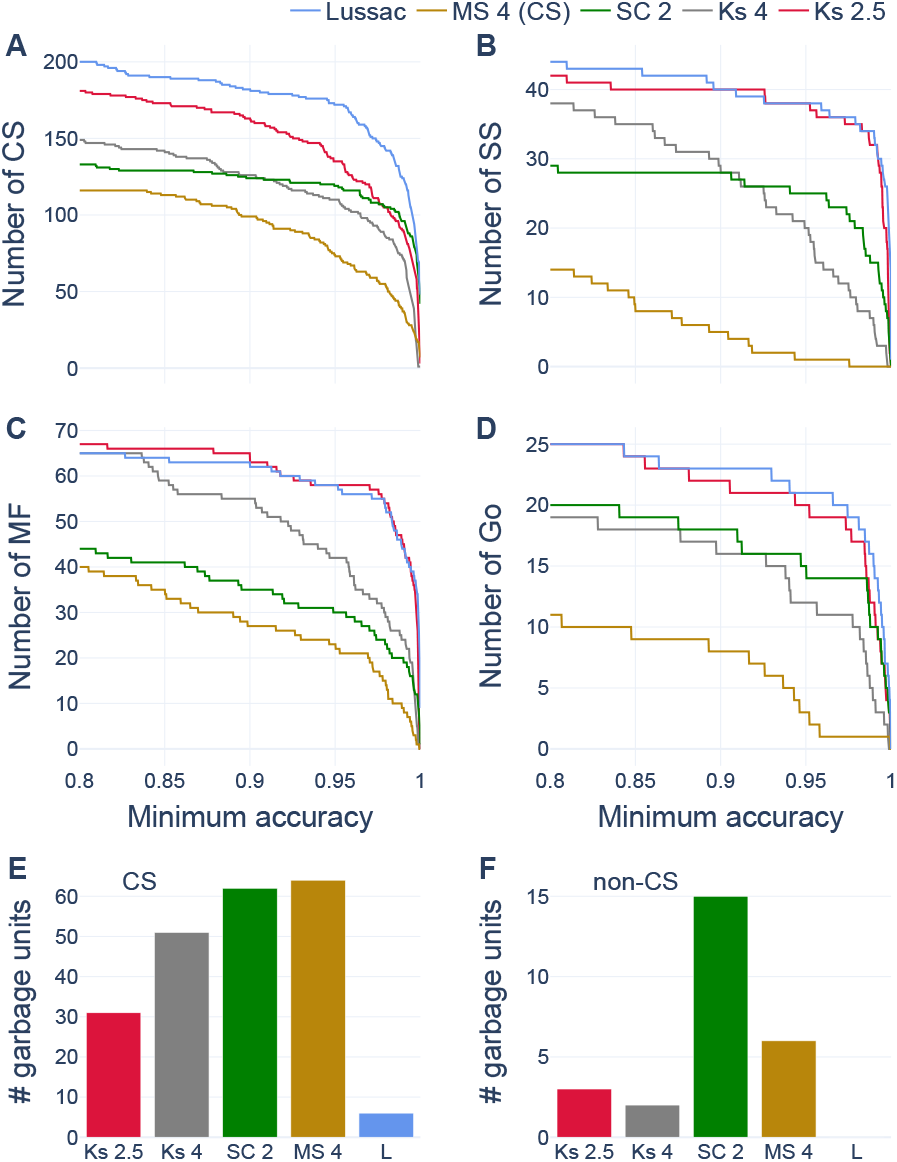
Cerebellar injection benchmark. Number of injected units recovered with an accuracy greater than or equal to the value on the *x*-axis for complex spikes (**A**), simple spikes (**B**), mossy fibers (**C**) and Golgi cells (**D**). (**E**) Number of garbage complex spikes units for each analysis. (**F**) Number of garbage non–complex units for each analysis. Garbage units are defined in Methods 3.4.6.

We also counted the number of garbage units, defined as units with low precision but relatively high recall (see Methods 3.4.6 for more details). As shown in Fig. 6E,F, *Lussac* significantly reduced the number of garbage complex spike units and output no garbage non–complex spike units.

#### 1.3.3 Computational time

A single run of *Lussac* is obviously computationally more expensive than running a single pacakge. Most of the difference arises from the need to run multiple analyses. The runtime for *Lussac* is added to this total but only represents about 10 to 40 % of the total (see Tables. 1, 2, 3 in the Methods). The whole process is moreover completely automated.

**Table 1.** Comparison of algorithms (synthetic datasets, stationary). def, default and opt, optimized parameters. l, light and h, heavy versions of *Lussac*.

| Analysis | Time (min) | $\geq 95\%$ | 80–95 % | 50–80 % | $< 50\%$ | Split | Red. | Merge | MUA | Noise |
| --- | --- | --- | --- | --- | --- | --- | --- | --- | --- | --- |
| KS 2 (def) | 22 | 195 | 58 | 23 | 21 | 8 | 17 | 7 | 22 | 5 |
| KS 2 (opt) | 33 | 233 | 56 | 34 | 20 | 8 | 7 | 12 | 19 | 5 |
| KS 2.5 (def) | 19 | 185 | 74 | 28 | 17 | 8 | 14 | 6 | 31 | 4 |
| KS 2.5 (opt) | 75 | 235 | 87 | 38 | 35 | 14 | 30 | 14 | 34 | 9 |
| KS 3 (def) | 37 | 117 | 59 | 40 | 27 | 22 | 5 | 11 | 9 | 1 |
| KS 3 (opt) | 49 | 119 | 66 | 15 | 2 | 22 | 26 | 2 | 0 | 1 |
| KS 4 (def) | 67 | 150 | 67 | 43 | 21 | 7 | 3 | 12 | 43 | 15 |
| KS 4 (opt) | 317 | 178 | 75 | 15 | 6 | 19 | 3 | 5 | 6 | 4 |
| SC 1 (def) | 113 | 109 | 55 | 47 | 32 | 26 | 0 | 23 | 14 | 0 |
| SC 2 (def) | 92 | 171 | 22 | 18 | 11 | 6 | 0 | 17 | 6 | 2 |
| TdC (def) | 192 | 39 | 41 | 9 | 0 | 0 | 0 | 6 | 0 | 0 |
| Total | 1016 | 251 | 77 | 37 | 26 | 0 | 0 | 0 | 0 | 0 |
| Lussac (l) | 63 | 253 | 68 | 25 | 16 | 0 | 0 | 9 | 9 | 1 |
| Lussac (h) | 145 | 249 | 73 | 30 | 13 | 0 | 2 | 10 | 14 | 0 |

**Table 2.** Comparison of algorithms (synthetic datasets, drifting). def, default, opt, optimized parameters. l, light and h, heavy versions of *Lussac*.

| Analysis | Time (min) | $\geq 95\%$ | 80–95 % | 50–80 % | $< 50\%$ | Split | Red. | Merge | MUA | Noise |
| --- | --- | --- | --- | --- | --- | --- | --- | --- | --- | --- |
| KS 2 (def) | 21 | 61 | 37 | 44 | 57 | 23 | 3 | 75 | 52 | 4 |
| KS 2 (opt) | 31 | 55 | 38 | 39 | 46 | 24 | 4 | 90 | 23 | 4 |
| KS 2.5 (def) | 20 | 48 | 64 | 68 | 108 | 96 | 6 | 14 | 63 | 6 |
| KS 2.5 (opt) | 78 | 45 | 63 | 132 | 108 | 176 | 12 | 28 | 57 | 15 |
| KS 3 (def) | 37 | 63 | 56 | 47 | 59 | 73 | 1 | 15 | 16 | 3 |
| KS 3 (opt) | 45 | 59 | 63 | 23 | 26 | 62 | 5 | 3 | 0 | 1 |
| KS 4 (def) | 68 | 96 | 78 | 58 | 37 | 50 | 1 | 26 | 37 | 7 |
| KS 4 (opt) | 329 | 113 | 62 | 35 | 26 | 74 | 3 | 13 | 11 | 14 |
| SC 1 (def) | 109 | 0 | 8 | 25 | 30 | 12 | 0 | 30 | 27 | 0 |
| SC 2 (def) | 99 | 84 | 20 | 23 | 18 | 19 | 0 | 9 | 9 | 5 |
| TdC (def) | 168 | 8 | 12 | 16 | 5 | 7 | 0 | 2 | 0 | 0 |
| Total | 1006 | 147 | 86 | 72 | 42 | 0 | 0 | 0 | 0 | 0 |
| Lussac (l) | 58 | 141 | 68 | 49 | 19 | 11 | 6 | 5 | 2 | 0 |
| Lussac (h) | 117 | 151 | 65 | 66 | 33 | 19 | 18 | 13 | 10 | 0 |

**Table 3.** Comparison of algorithms (injection datasets). Comparing the total run time of each individual analyses, of all of them combined (total), and of the *Lussac* combination. Also comparing the amount of injected cells recovered with an accuracy greater than 80 %.

| Analysis | Run time (min) | CS (/239) | SS (/53) | MF (/86) | Go (/32) | MLI (/12) | PLI (/4) |
| --- | --- | --- | --- | --- | --- | --- | --- |
| KS 3 | 84 | 116 | 20 | 29 | 11 | 4 | 0 |
| KS 4 | 82 | 149 | 38 | 65 | 19 | 6 | 0 |
| SC 2 | 118 | 133 | 29 | 44 | 20 | 6 | 0 |
| KS 2 SS | 121 | 168 | 42 | 67 | 25 | 8 | 0 |
| KS 2 CS | 124 | 181 | 40 | 60 | 24 | 8 | 0 |
| KS 2.5 op | 111 | 54 | 42 | 67 | 25 | 8 | 0 |
| MS 3 CS | 79 | 127 | 1 | 17 | 4 | 0 | 0 |
| MS 4 CS | 281 | 116 | 14 | 40 | 11 | 1 | 0 |
| Total | 1000 | 209 | 44 | 76 | 26 | 8 | 0 |
| Lussac | 328 | 200 | 44 | 65 | 25 | 8 | 0 |

### 1.4 Limitations

Although we have been able to develop a robust framework for generating a consensus output that improves upon individual packages under most conditions, *Lussac* is not perfectly robust or free from bias.

Spike sorting is a noisy analysis and some apparently high-quality but incorrect units will be generated by chance. Because *Lussac* combines many analyses, there can an increased risk of some such incorrect units being retained in the final consensus output. Avoiding this requires balancing the stringency of the consensus-forming mechanisms against the yield. We believe this has been achieved with success, as detailed above, but it is important to recognize that the mechanism exists and it is likely that fine-tuning of the decision thresholds internal to *Lussac* could further improve performance.

The difficulty of correctly excluding apparently high-quality but incorrect units increases if a package generates large numbers of them. In our experience, this was the case for MountainSort (probably due to the poor recovery of overlapping spikes), leading us to exclude it from use in *Lussac* except for the special case of finding complex spikes in cerebellar recordings, in which its benefits outweighed any problems. This phenomenon is unproblematic as long as good analyses in a reseanable number are used. Recommendations about the packages to include are included in the detailed online documentation.

Despite using multiple approaches to benchmark package performance, there remain known issues. The spike waveforms in MEARec are unusually long-lasting and this may induce a reduction in performance for packages with default parameters. The spike injection procedure did not respect the geometric orientation of cells.

## 2 Discussion

We have developed and validated a consensus spike-sorting package able automatically to exploit the information from multiple different analyses. The foundations of our package involve the development, extension and formalization of several metrics for evaluating unit quality and similarity. Thus, although violations of the refractory period represent a classical measure of unit quality, we have provided a more complete analysis for its quantitative interpretation than was available in the prior literature. The significant innovations in metrics emerge from the processing of units from different analyses, because *Lussac* is the first package to automate such a consensus procedure. The key novel metric is the cross-contamination, which is integral to detecting erroneous merges. Our multi-analysis approach showed that these merges are problematic for most spike-sorting packages applied individually.

The core steps of our algorithm involve the creation, refinement and analysis of a graph incorporating the outputs of all of the individual analyses. The graph’s nodes (units) and edges are parameterized with the complete set of unit quality and comparison metrics. We have developed a multistep procedure for analyzing this graph, in particular to identify erroneous merges. Important subsequent steps of our analysis involve the selection of the best unit for each neuron and, if beneficial, the construction of a consensus spike train. Each of these steps is guided by quantitative quality metrics.

Our benchmarking shows that *Lussac* can benefit spike-sorting analyses in several ways. Firstly, a greater number of correct units can be detected, because individual packages nearly always miss some. Secondly, *Lussac* generally improves the quality of the units detected, because better representations can be constructed from the multiple analyses. Thirdly, the package leverages the multiple analyses to eliminate a substantial proportion of erroneous merges, a problem that is essentially insoluble within individual analyses. Thus, *Lussac* avoids to a significant extent the trade-off between false negatives and false positives that the individual packages face. It is worth pointing out that a simplistic comparison of spike sorters based upon the number of units reported would penalize the removal of incorrect units. It is therefore critical to benchmark packages using some form of ground truth.

For analyses with clean synthetic data, *Lussac* performs at least as well as each of the individual packages tested. In more challenging situations involving electrode drift or more varied spike forms, notably including complex spikes, the benefits of *Lussac* over individual packages are more pronounced. These tests demonstrate the difficulty for individual packages to perform well under all conditions and highlight the compromises most have to make.

*Lussac* increases the yield and quality of the data, but also eliminates manual post-processing and, less obviously, the effort that would have been spent optimizing the choice of package and parameters. A further benefit of a consensus method is that it limits the bias towards specific neuronal types inherent in the use of a single package and a single parameter set. Thus, despite the increased computational resources required to operate *Lussac*, operator productivity is improved.

Lussac version 2.0 is publicly available under a free-software licence. It relies upon and has been partially integrated into the SpikeInterface package. It can be operated as a practical analysis package but may also serve as proof of principle encouraging other spike-sorting packages to incorporate the metrics and consensus methods we have developed. Thus, our procedure for intra-analysis unit merges has already been ported to the development version (2) of SPYKingCIRCUS and portions of the work underlying *Lussac* have been contributed to SpikeInterface.

Amongst possible directions for future progress, it is clear that the multi-analyses graph could be analyzed using many different approaches and it is probable that further improvements in spike sorting will result. A more comprehensive analysis (e.g. analyzing the graph as a whole instead of individual connections) to produce the consensus would likely facilitate fine-tuning of the tradeoff between stringency and yield that influences the robustness of *Lussac* when faced with inputs containing many low-quality units.

## 3 Methods

The *Lussac*^1^ software, developed in Python3, is available under a free-software license on GitHub (https://github.com/BarbourLab/lussac). It relies on SpikeInterface [10], into which we have integrated several sections of code.

### 3.1 Signal filtering

In several steps of the pipeline, *Lussac* needs to extract traces from the recording. To reduce the impact of noise, filtering is crucial.

Every time-domain filter in *Lussac* is implemented with a Gaussian filter. This is a time-symmetrical filter that minimizes rise and fall times, while having the property of not overshooting (nor therefore ringing) in response to a step input. As a Gaussian filter is only a low-pass filter, we implemented the high-pass filter by subtracting the filtered trace from the original trace. In this way, a bandpass filter can also be generated with any desired cutoff frequencies.

### 3.2 Unit metrics

#### 3.2.1 Signal-to-Noise Ratio (SNR)

Units with a higher signal-to-noise ratio tend to be more reliable, as they are easier to detect and cluster. However, when comparing units, a higher SNR is not necessarily better, as it might result from reduced detection of spikes with lower amplitude.

We defined the signal as the peak absolute voltage of the template across all channels, and the noise as the baseline standard deviation (estimated from the median absolute deviation) on the same channel. It is important to note that the SNR is dependant on the filter used.

#### 3.2.2 Spike number/frequency

In general, if a unit recovers more spikes, we consider it to be of higher quality. However, this does not distinguish between recovery of true spikes and contaminant spikes. Thus, we chose to use this metric in conjunction with the contamination metric (see Results 1.1.1).

Different neuronal types are of course expected to have very different firing rates, so this metric is best applied in a relative manner, when comparing units representing the same neuron.

The total number of spikes in a unit is also important in governing the precision of the other metrics, notably those based upon correlogram comparisons and refractory period violations. With low numbers of spikes, some of the metrics become less accurate, and may not be usable in practice. However, in most cases, a sufficient number of spikes can be ensured by having a long-enough recording.

#### 3.2.3 Contamination ratio

The classical absolute refractory period (i.e. the time during which a second action potential is impossible, irrespective of the drive) can be shorter than a millisecond at physiological temperature. However, in some neurons, longer intervals are in practice rarely invaded, presumably because insufficient excitatory drive is available during this *apparent* refractory period. Thus, we shall often count violations in this apparent refractory period for higher accuracy. In an abuse of terminology, we shall continue to call this the refractory period throughout this paper.

We censored a small window following each spike, because detection of spikes within this window is often unreliable and their processing is inconsistent between packages. The width of this window is typically *t*_*c*_ = 0.4 ms (but can be adjusted depending upon the spike width). Hence, in equations 1 and 2:

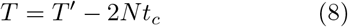

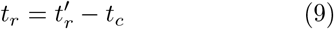

where *T*^′^ and 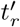 are the real durations of the recording and refractory period, and *N* the total number of spikes. These corrections are not mathematically exact, as the correction actually depends on the ISI (Inter-Spike Interval) distribution, which is unknown. They are however close enough for a good approximation.

A mathematical proof of equations 1 and 2 is given in Appendix G. These equations may not have a real solution if *n*_*v*_ is large, which can arise in practice if the contaminant spikes are strongly correlated, thus causing a departure from the assumption of independant firing. In such cases, we simply follow the convention of setting *C* = 1.

In our method, we decided to choose equation 2 because we consider it more plausible. If the assumption of randomness is not satisfied, this choice can lead to a slight underestimation of the contamination. For example, if *C*_∞_ = 0.1, then *C*_1_ ≈ 0.106.

It should be noted that the contamination ratio is an expectation about which actual values will vary, because of the variability of *n*_*v*_. The coefficient of variation (sd/mean) of the contamination ratio can be approximated by 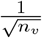, which, for a given contamination, decreases as *t*_*r*_, *T* and the mean firing frequency increase. This metric will therefore work better with longer recordings.

Both positive and negative correlations between neurons can occur within the refractory period. This leads to an error in the estimated contamination, in the sense that a positive correlation causes an overestimate of contamination and an anti-correlation causes an underestimate. This can occur during oscillations or because of the network connectivity. However, more important than the absolute value of the contamination ratio for our procedure are the relative values when comparing units representing the same neuron. Apparent anti-correlation between neurons can also arise artefactually, as a result of poor or incomplete treatment of spike collisions in spike-sorting packages, which is the main reason why the censored period is important.

Eq. 1 is equivalent to the result obtained in [11], except that they evaluated refractory period violations of the ISI distribution, which is mathematically inaccurate (e.g. if three spikes are detected in a very short period of time, then the violation between spikes 1 and 3 should also be counted). Furthermore, they made the hypothesis that random “rogue spikes” only interact with true spikes, which seems more consistent with the case of a single neuron with its own refractory period being responsible for the contaminating spikes.

#### 3.2.4 Waveform variability (SD ratio)

The hypothesis of this metric is that invariant spike amplitudes are measured after the addition of noise. The standard deviation of the measured amplitudes should therefore be equal to that of the background noise. A deviation from this equality, usually an increase of standard deviation of spike amplitudes, indicates a problem in the unit, typically contamination.

To reduce the impact of bursts on the standard deviation of spike amplitudes, we added a censored period (4.0 ms by default), whereby spikes following other spikes within this censored period are eliminated from the calculation. This prevents spikes that are usually of lower amplitude from impacting the standard deviation.

Electrode drift can also strongly influence the standard deviation, because spike amplitudes on a given electrode will change. To minimise the contribution of drift-related spike amplitude variations, we subtracted successive spike amplitudes. This isolates the typical spike-to-spike variations of measured amplitudes while minimizing the influence of slow or infrequent rapid drift events. Under the hypothesis that successive spike amplitudes are independent, this leads to a distribution of differences whose variance is double that of the noise added to the spikes. We thus divide by 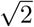 to recover an SD estimate that is less sensitive to drift.

Finally, the presence of the spike itself contributes to the standard deviation of the noise computed on the same channel. This noise source may be non-negligible for units with a high firing rate and a high amplitude. To compute the SD of background noise, a simple approach would be to subtract the template for the spike from the trace at all the times the spike was detected and then to measure the SD. However, this is computationally costly, because the procedure must be repeated for every unit. Instead, we decided to approximate the calculation mathematically. Under the assumption that the templates and noise are two independent random variables (which is not exact, as it does not take into account their time correlation), their variances sum. We can thus estimate the SD of the noise in the absence of the the spikes of the unit under consideration:

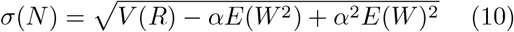

where *E, V* stand for the expected value and variance respectively, *N* is the underlying noise, *R* the voltage trace, and *W* the template. *α* stands for the fraction of the recording the templates span. See Appendix F for the detailed calculation.

Improvements could also be envisaged but have not been implemented to correct for the sampling jitter, which can affect both the amplitude and SD of a spike waveform [12].

#### 3.2.5 Generating a unit quality score

A unit is composed of correctly detected spikes (True Positives; *TP*), of contaminant spikes (False Positives; *FP*) and may be missing some spikes from the neuron (False Negatives; *FN*). The quality metric in equation 3 can be expressed using these values. The number of spikes in the unit is thus *N* = *TP* + *FP*, and the contamination ratio 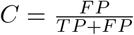. Additionally, the number of spikes in the neuron is *N*^′^ = *TP* + *FN*, where both *N*^′^ and *FN* are unknown. Given this, we can calculate:

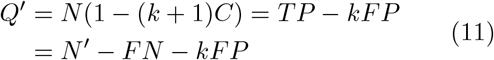

In this view, *k* allows weighting of *FP* compared to *FN*. When comparing two units representing the same neuron, the quality will be increased if:

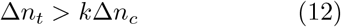

where Δ*n*_*t*_ and Δ*n*_*c*_ are differences in *TP* and *FP* between both units.

#### 3.2.6 Spike coincidence

When comparing multiple analyses, the spike trains of units representing the same neuron will usually have numerous simultaneous spikes. Detection of coincident spikes is relatively straightforward: we count the number of spikes *n*_*s*_ in one unit occurring within a certain window *w* (± 0.3 ms by default) of a spike in the second unit. However, coincident spikes can occur by chance between two uncorrelated units, with a probability 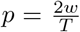 for each pair of spikes. Using a reasoning similar to that in Methods 3.4.3, we can correct for this and compute the “real” number of coincident spikes from spikes representing the same neuron:

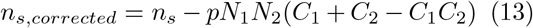

where *N*_1_, *N*_2_ are the number of spikes in each unit, of respective estimated contamination *C*_1_, *C*_2_. From this, we can compute a simple similarity metric:

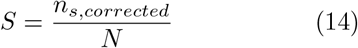

A value of *S* = 0 is expected between two uncorrelated neurons, while a value of *S* = 1 means a perfect correlation. A negative value indicates an anti-correlation.

This metric could be constructed directionally, depending on the value of *N* that is used: *N* could take the value of *N*_1_ or *N*_2_. However, we chose a single bi-directional metric that combines all the information from both units, using *N* = min(*N*_1_, *N*_2_). This choice was made so that if all spikes from a unit are included in the spikes of another unit, then the similarity becomes 100 %, allowing small split units to have a high similarity.

#### 3.2.7 Template comparison

A fundamental metric for comparing two units is the similarity of spike waveforms. The unit templates were computed as the averages of up to 2,000 unique waveforms on the 5 common channels with the largest absolute signals. Signals were bandpass-filtered with a Gaussian kernel (150–7000 Hz by default).

For this metric to be reliable, it is important to correct any misalignment. In the inter-analysis merging step, this is done using the cross-correlogram (see section 1.2). However, for the intra-analysis step, units seldom have coincident spikes. We thus computed the template similarity metric in Eq. 5 for all possible lags between the units in a window of ± 0.17 ms and kept the lag with the lowest value for the metric.

**Supplementary Fig. 1.**
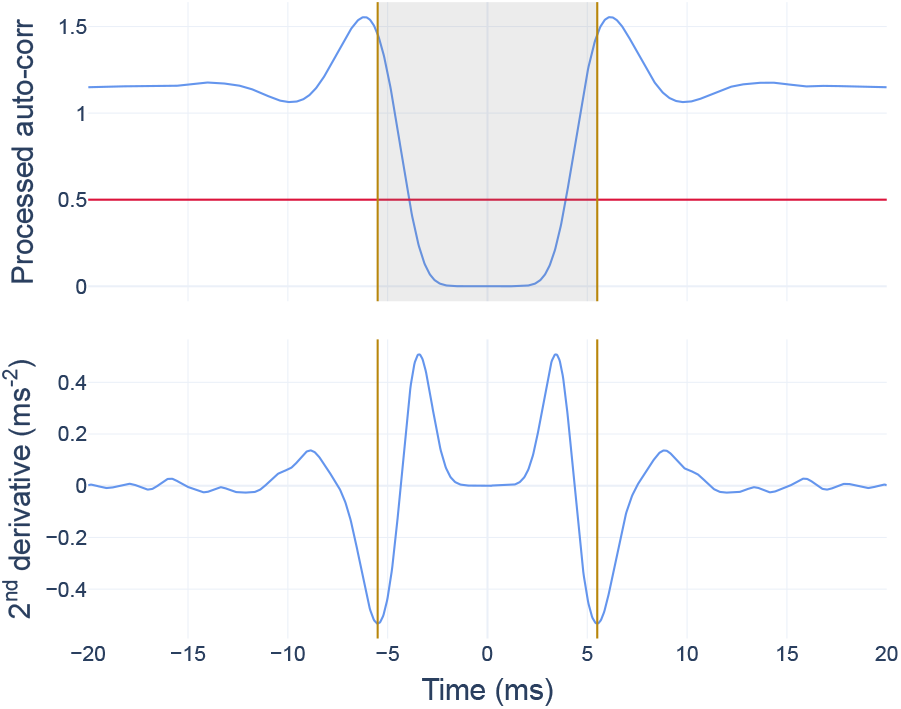
Determination of auto-correlogram comparison window. (**Top**) Normalized filtered auto-correlogram. (**Bottom**) Second derivative of the correlogram above. The gold vertical lines indicate the most central minima of the second derivative where the correlogram value simultaneously exceeds half its mean (red horizontal line). These minima delimit the window for correlogram comparisons (gray shaded area).

#### 3.2.8 Correlogram comparison

It is expected that units representing the same neuron will have similar auto-correlograms and that these resemble the cross-correlogram between them. However, as we formalize below, these similarities are, under randomness assumptions, statistically exact and extend to daugther units of a split parent even under extreme sampling regimes. Comparisons of correlograms therefore constitute a powerful metric for detecting split units and more generally those representing the same neuron.

To mitigate the noise due to binning before comparing correlograms, they are Gaussian-filtered with a standard deviation of 0.6 ms. Because some packages detect the same spikes multiple times, it is important then to censor (i.e. set to zero) the period between ± 0.3 ms.

In general, only the region of the correlogram around zero lag (the central dip) is useful for comparisons. The correlogram at larger shifts is often featureless, including them introduces noise into the comparisons. We therefore extract the “window” of each correlogram according to a procedure that adapts robustly to the varied correlogram shapes of different units, as shown in Fig. S1. The second derivative is computed, and the most central minimum that exceeds half of the average correlogram value is taken. For each comparison between two units, the window for comparison is given by *w*, the weighted sum:

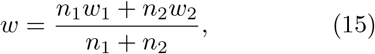

where *w*_1_, *w*_2_ are the windows for each auto-correlogram, and *n*_1_, *n*_2_ are the numbers of spikes in each unit.

Given an auto-correlogram *A* and a cross-correlogram *C*, we compute the average absolute difference:

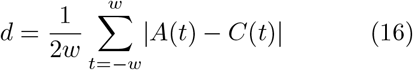

#### 3.2.9 Cross-contamination

A central issue when refining the multi-analysis graph is the identification of merges that create an incorrect link between units representing different neurons (the “triplets”). Such triplets can be merges or splits. A key difference between these two cases is that the spikes of a merge can violate the refractory periods of spikes in two units in another analysis, because each represents a single neuron. We developed a metric to quantify this cross-contamination. The metric has the downside of only looking at the proportion of spikes in unit *B* not coming from the same neuron as the one represented by unit *A*, meaning that it does not distinguish between random contamination in unit *B* and the possibility that it represents another neuron. More complex models were explored to make this distinction, but we found that they were not robust and gave absurd results very quickly when the initial assumptions (such as independence of activity) were not met. As these assumptions are in reality often not met completely, we chose to use the simpler and more robust model.

Under this model, the expected number of cross-violations *n*_*cv*_ can be well approximated by a binomial distribution: for each pair of spikes (one spike from each unit), there is a probability 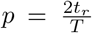 that the spike pair will violate the refractory period. The total number of spike pairs that can give rise to violations is given by: *n*_*cv*_ = *n*_*A*_*n*_*B*_(*C*_*A*_ + *CC*_*B*→*A*_ − *C*_*A*_*CC*_*B*→*A*_), where *C*_*A*_ is the contamination of unit A, and *CC*_*B*→*A*_ is the cross-contamination between units A and B, with A the reference. Thus, *n*_*cv*_ ∼ ℬ (*N, p*).

For a given threshold *X* on the cross-contamination, we can compute the probability of observing *n*_*cv*_ by setting *CC*_*B*→*A*_ = *X*. Thus:

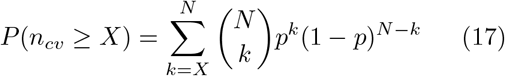

As mentioned in the Results section, this metric is directional. However, unless specified otherwise, we generally take as reference (A) the unit with the lowest estimated contamination.

### 3.3 *Lussac* pipeline

Having only briefly described the merge modules in the Results, we now describe in depth all of the different modules available in Lussac, and supply additional information about both merge modules.

#### 3.3.1 Unit alignment

A necessary preliminary step before any comparisons between units is to ensure, as far as possible, a canonical alignment. It is not possible to guarantee a unique aligment, as slight changes in the average waveform can occur, so we adopted several heuristics with the aim of creating a procedure that is robust. We first computed the filtered templates on the best channel (200– 6000 Hz by default), and identified the first peak greater than half of the peak absolute value over the whole template (preventing the alignment of multi-phasic waveforms from depending on which peak had the highest amplitude). We then performed a local search for a greater maximum over the following 0.33 ms to reduce sensitivity to noise. By default, this whole procedure is performed on ± 2 ms, but asymmetrical windows could be beneficial in certain cases, notably for cerebellar complex spikes.

#### 3.3.2 Unit annotation

It can be useful to identify types of neurons recorded, as post-processing might be different for different types. Many sources of information can be exploited to identify specific neuronal types, such as the mean firing rate, estimated contamination, Signal-to-Noise Ratio (SNR), SD ratio, ISI fraction. For each, the user can provide lower/upper limits. The “ISI fraction” configuration parameter represents the fraction of spikes that have inter-spike intervals falling in a given range. Additional sources of information for annotation might include spatial coordinates and interactions between neurons revealed by cross-correlograms.

At this time in Lussac, a concrete implementation only exists to identify cerebellar Purkinje cell complex spikes, as their processing is quite different. We classed as putative complex spikes those units with a firing rate between 0.2 and 5 Hz, and a low proportion (*<*5 %) of spikes with 10.0 *< ISI <* 35.0 ms (“ISI fraction”). A more general annotation package for cerebellar neurons has been developed [14].

#### 3.3.3 Removal of duplicate spikes

Some spike-sorting algorithms detect the same spike multiple times. In order for metrics to work as intended, it is beneficial to remove such duplicate spikes. It is also necessary as part of the merging procedures. The default behaviour is to scan spikes of a unit and to remove succeding spikes with intervals smaller than 0.3 ms.

#### 3.3.4 Removal of redundant units

Redundant units (i.e. units that share > 70 % spikes in a ± 0.3 ms window) are eliminated. Different criteria exist for choosing which unit to keep, but the default behaviour is to retain that with the largest amplitude in its template.

#### 3.3.5 Removal of low-quality units

Before combining multiple analyses, it can be beneficial to remove units that are clearly of low quality. This can accelerate the subsequent analyses, and reduce the probability of wrongly connecting units in the graph analysis. The user can configure this in the exact same way as described in Unit annotation, for instance by removing units with unreasonably low firing rates, which would compromise the use of the SD-ratio metric and correlogram comparisons.

#### 3.3.6 Intra-analysis unit merge

A number of sanity checks and quality criteria must first be satisfied by candidate units: they must have a sufficient number of spikes to enable accurate correlogram comparisons (1,000 by default) and their estimated contamination must be less than 20 % (to enable comparison of the central dip).

For all pairs of units, we calculate the differences of their waveforms Δ_*W*_ and correlograms Δ_*c*_ (described above). The units of pairs with Δ_*W*_ *<* 0.25 and Δ_*c*_ *<* 0.16 are considered to come from the same neuron. In order to filter out the worst units for the merge step to follow, we compute the quality score of Eq. 3 (without SD ratio for faster computation) of each unit of the pair and also of the putative merge (which requires the removal of duplicated spikes, as described above). If the merge would result in a unit of higher score, both units are retained unmerged (merging happens at the next step). Otherwise, the unit with lowest quality is removed.

Groups of units are then formed, where each group contains all units considered to represent the same neuron according to the criteria of the last paragraph. Membership of a group can be indirect. Thus, if two pairs A-B and B-C are retained, then units A, B and C are considered to represent the same neuron. For each such group, an iterative merge process is then carried out as follows. Pairwise trial merges, where spike trains are combined and duplicates removed, are performed for all pairs in a group. The merge with the highest score is retained and then the next iteration begins. This process stops once a single node remains or the quality of a individual node is higher than that of any possible merge, in which case that node is retained.

#### 3.3.7 Inter-analysis unit merge

##### Pairwise alignment

The cross-correlogram between units of all analyses are computed on a window of ± 1.33 ms (by default, the maximum shift possible) and convolved with a Gaussian kernel of standard deviation 50 µs, as diagrammed in Fig. 1A. Then, a threshold is set to 5 % of the number of spikes in the smallest unit. If a peak in the cross-correlogram crosses this threshold, then the timing of the peak corresponds to this shift. Otherwise, the shift is set to zero.

##### Removal of merged units

This stage focuses on triplets of units where a unit from one analysis is connected to two units from another analysis, as diagrammed in Fig. 3D.

To ensure robust decisions in the presence of variability resulting from low spike counts, the cross-contamination criterion, which is critical for detecting merged units, is implemented as a statistical test (see 3.2.9). This reports the probability of observing the number of cross-violations seen between both units, if the cross-contamination is at least 10 %.

If the p-value exceeds 5 × 10^−3^, the triplet is considered to contain a split rather than a merge and the algortihm moves on to the next “triplet”. Conversely, if *p <* 5 × 10^−3^, then we consider whether the node central to the “triplet” is a merge by comparing it to the other two nodes. If both its contamination and SD ratio are higher than those of the other two units, then the central node is removed. Additionally, if the p-value is really low (*p <* 10^−8^), the central node is removed irrespective of the value of the SD ratio.

If the central node is not removed in this way, then we examine whether one of the peripheral nodes is a garbage unit accounting for the high cross-contamination. We compute the cross-contamination between the central node and each peripheral node (taking the central node as reference), and if the p-value is less than 5 × 10^−3^ and the peripheral node has a higher contamination, then the peripheral node is removed. Otherwise, no node is removed.

##### Removal of bad connections

When checking every edge of the inter-analysis graph, all the inter-unit metrics are used. There are two cases where the edge is considered problematic:

| Similarity | Cross-cont. (6 %) | $\Delta_T$ | $\Delta_c$ |
| --- | --- | --- | --- |
| X | $p < 5 \times 10^{-3}$ | $> 0.10$ | $> 0.06$ |
| $< 0.60$ | $p < 0.80$ | $> 0.05$ | $> 0.05$ |

where the cross-contamination test is performed with a reference value of 6 %, and $\Delta_T, \Delta_c$ are the template and correlogram differences. If either of these cases apply, then the edge is removed.

Furthermore, removal of the edge suggests that it may be beneficial to remove at least one of the nodes. For this reason, a check is performed on the normalized differences in contamination and SD ratio between both nodes. Thus, a node *A* is removed if both following conditions are satisfied:

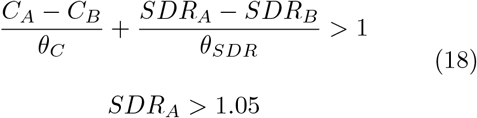

where *SDR* is the SD Ratio and *C* the contamination. The contamination and SDR differences are normalized by their respective thresholds with values *θ*_*C*_ = 0.08 for the contamination and *θ*_*SDR*_ = 0.20 for the SD ratio. The first condition represents a diagonal line in the contamination– SDR plane, such that a node is removed if its normalized contamination difference is high, if the normalized SDR difference is high or if a combination of the two is high, subject to the additional condition that the SD ratio of node *A* must also exceed 1.05. Thus, if one of the nodes connected by a removed edge is clearly of lower quality than the other, the worse node is in turn removed.

### 3.4 Benchmarks

#### 3.4.1 Creation of a synthetic neocortical dataset

A wholly synthetic dataset was created using the MEArec package [13]. The recording was made simulating 128 channesl with a Neuropixels 1.0 layout. The 3,000 neurons were distributed randomly by MEArec in a box of size 152 × 320 × 1320 µm, giving a density of 46 725 neurons*/*mm^3^. Neurons were uncorrelated, non-bursting and spaced at least 18 µm center-to-center. The voltage traces were simulated for 1 hour. 80 % of the neurons were excitatory (with a mean firing rate of 5 Hz), the rest were inhibitory (with a mean firing rate of 15 Hz). An additive white noise was generated with a Gaussian process of standard deviation 10 µV.

For the dataset with drift, rigid^2^ drift was implemented. Fast drift events happened every 10 minutes, with an amplitude of 5 to 20 µm. Slow drift was implemented as a sinusoid of 55 µm amplitude peak-to-peak, and a maximum velocity of 2 µm min^−1^.

#### 3.4.2 Automated post-processing

Before comparing each analysis, we removed the obviously garbage units. The rationale for this is that those units are very easy to remove and would pollute and slow the analysis (as they can be numerous). These obviously bad units were defined as those with an average firing rate lower than 0.4 Hz and/or an estimated contamination higher than 25 %.

#### 3.4.2 Comparison to a ground truth

##### Metrics

In comparing the output of spike-sorting packages with an exhaustive ground truth, a number of metrics combining true positives *TP* and true/false negatives (*TN, FN*) can be of interest:

- **Accuracy** 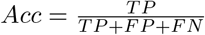 reports how well the neuron is represented by the unit, penalizing false positives and false negatives equally.
- **Recall** 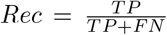reports the fraction of spikes from the neuron that were recovered by the unit.
- **Precision** 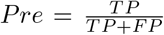reports the fraction of spikes from the unit belonging to the neuron.

Given no correlation between a neuron and a unit, some collision events are still expected to occur at random, with a probability 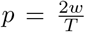 for each pair of spikes, where *w* represents the time window for two spikes to be considered as colliding (in our case *w* ≈ 0.27 ms), and *T* is the duration of the recording. A correction can be applied to take these random collisions into consideration. For a unit with *N* = *TP* + *FP* spikes, the expected number of collision events is:

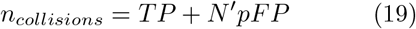

where *N*^′^ is the number of spikes in the reference neuron. From this formula, we can derive a novel estimate of the number of true positives:

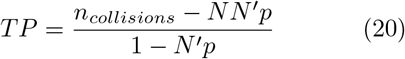

##### Automated categorization

Thanks to the exhaustiveness of the ground truth, it is possible to categorize automatically every unit in the output of a spike-sorting analysis. Our method works sequentially, as presented in Fig. S2. First, for each unit, the sum of precisions with all neurons is computed, which estimates the fraction of detected events that actually match a neuronal spike. If this sum is lower than 50 %, the unit is categorized as “Noise”. Then, for each uncategorized unit, if the value of the second highest recall if higher than 1/3 (meaning the unit recovered more than a third of the spikes from two distinct neurons), then it is categorized as “Merged”. Then, if the highest precision is lower than 50 %, it is categorized as “MUA” (Multi-Unit Activity), because more than half the spikes don’t come from a single neuron. At this stage, uncategorized units are considered good enough to be matched to a neuron (however, one neuron can still match to multiple units). For each neuron, the uncategorized unit with the highest accuracy is considered “Matched”, while the others are either “Redundant” (if the fraction of overlap with other units is higher than 50 %) or “Split”.

#### 3.4.4. Real cerebellar cortical recordings

Inclusion of the MountainSort 3 and 4 packages improved consensus performance for the recovery of cerebellar complex spikes. By default, the voltage-time snippets excised and analyzed are symmetrical around detected spike. For complex spikes, a long window must be set to include the whole waveform, but because of the symmetry, almost half of the snippet, before the complex spike, contains no useful information and injects noise into the analysis. We therefore patched MountainSort 4 to implement an asymmetrical snippet, with a short time before and a longer time after the detection of the complex spike. A git pull request has been created for this patch: https://github.com/llobetv/ml_ms4alg

The acquisition of this dataset is described in detail in the corresponding paper (Llobet et al, in preparation), and the data will be made available when this paper is published. In brief, 4- or 6-shank, 64-site multi-electrodes (Cambridge Neurotech) were implanted in mouse cerebella and advanced towards the putative eyeblink region until Purkinje cells responding strongly to the air puff directed at the ipsilateral eye were recorded. Continuous recordings throughout learning and extinction sessions (3 sessions of 30 minutes per day and 75 trials per session) were obtained. The mice were walking continuously on a motorized wheel; the possible influence of the locomotion on the cerebellar activity should be borne in mind. The recordings were band-pass filtered at acquisition (0.1–7500 Hz).

#### 3.4.5 Linking Purkinje cell simple and complex spikes

The simple spike–complex spike duality of the Purkinje cell allows their unequivocal identification and constitutes a form of ground truth in real data. If the spikes come from the same Purkinje cell, then their filtered templates (*>*1500 Hz) are near-identical (see Appendix E). Furthermore, the cross-correlogram between them shows a ∼ 10 ms pause of simple spikes after the occurence of a complex spike. If we find both of these features, then we can be almost certain that these two units represent the simple and complex spikes of the same Purkinje cell.

#### 3.4.6 Spike injection

Cerebellar cortical recordings from eight different mice (coming from Real cerebellar cortical recordings) were analyzed with different spike-sorting algorithms (Kilosort 2 & 3, MountainSort 3 & 4 and SpyKING CIRCUS). High-quality units were selected manually and categorized as complex spike, simple spike, Golgi cell, mossy fiber, MLI (Molecular Layer Interneuron) or PLI (Purkinje cell Layer Interneuron) [15]. From each category, a set of templates was constructed: if a neuron was detected by multiple analyses, the best unit was chosen (using the quality metric of Eq. 3). The template was extracted from the filtered recording (60–12 000 Hz) with a window of − 15 to 25 ms around the peak. The borders were forced to go to 0 µV using MEArec tools [13]. Only channels with at least 40 µV of signal and that were at most 150 µm from the peak channel were kept. For each channel, the layer of cerebellar cortex the unit was in was annotated, to enable injection in the same layer in recordings from other mice.

**Supplementary Fig. 2.**
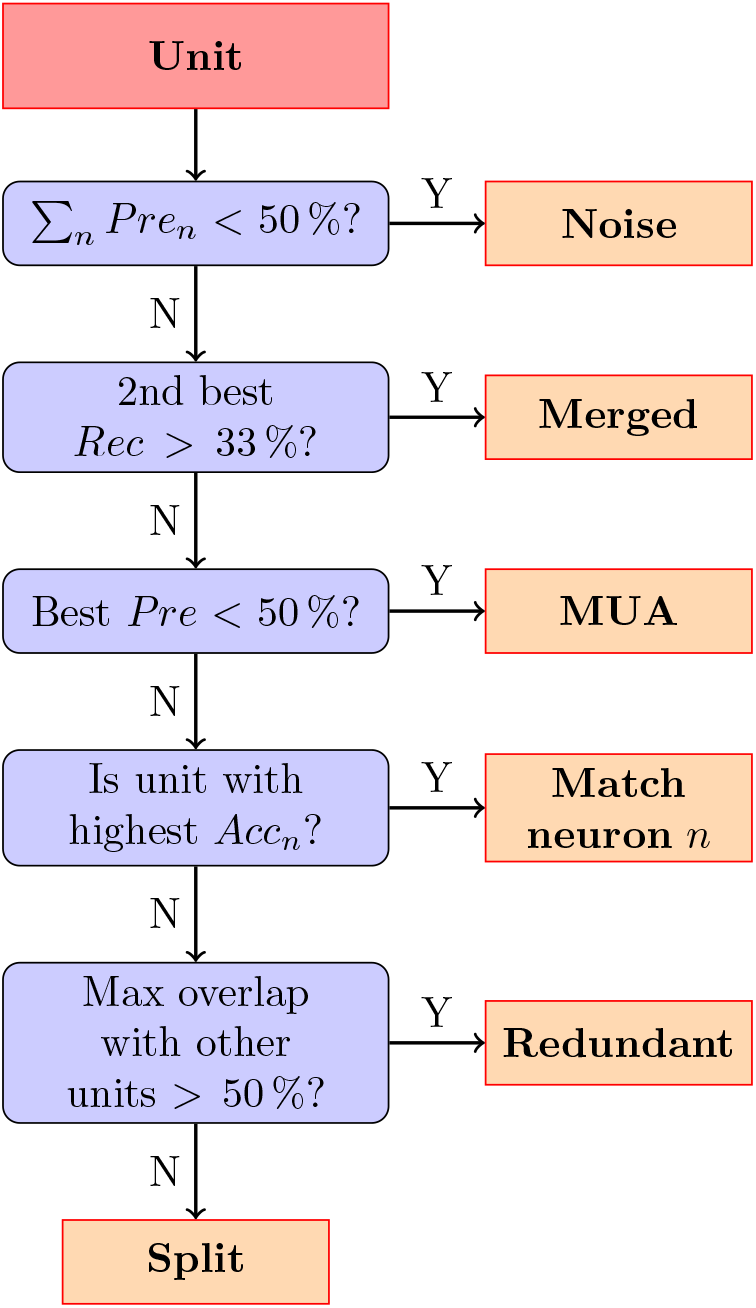
Scheme for unit categorization. *Pre*_*n*_ stands for the precision with regards to neuron with index *n*. Recall *Rec* and accuracy *Acc* behave similarly.

For each recording, all templates coming from other mice are considered individually. For each template, an injection location where all the channels were in the correct layer was sought. If multiple locations existed, one was chosen at random. If no such location exists, then a location with exactly one channel mismatch is accepted. If no such location existed, then the template was not injected. The timing of the original spike train was retained. If the recording was longer than the original one, then spikes were appended by sampling from the ISI distribution. The injected template was a scaled down version (sampled randomly within a range), to make it harder to detect.

This hybrid dataset does not contain an exhaustive ground truth. Therefore, we detected garbage units as those having a precision lower than 0.5 and a recall higher than 0.3 with at least one of the injected template. This means that a non-negligible portion of the injected neuron was recovered, but that the unit lacks the specifity to be considered a good unit.

### 3.5 Code and data availibility

The *Lussac* package, documentation and tutorials are publicly available under the AGPL3 licence (https://github.com/BarbourLab/lussac). The synthetic neo-cortical data, injection data and code used to generate the figures will be made available upon publication. For Fig. 5, the complete data set will be made available when the first paper analyzing it is published.

## Acknowledgements

We thank Jonas Ranft for useful discussions throughout the development of *Lussac*. We thank Idriss Tsayem, Guillaume Dugué and Brandon Stell for being early adopters of our algorithm and helping us with debugging. We also thank the whole SpikeInterface team, particularly Alessio Buccino and Samuel Garcia, for their precious help with the integration in the SpikeInterface framework.

This work was supported by the Fondation pour la Recherche Médicale (Equipe FRM DEQ20160334927), The Agence Nationale de la Recherche (ANR-22-CE37-0001-01) and the program ‘Investissements d’Avenir’ from the French Government, implemented by Agence Nationale de la Recherche (ANR), references: ANR-10-LABX-54 MEMOLIFE and ANR-11-IDEX-0001-02 PSL* Research University.

## Author contributions

Victor Llobet developed the contamination model, the unit quality metric (3), the initial correlogram comparison, the spike alignment method and patched MountainSort. He produced the initial version of the consensus algorithm, applied it to real cerebellar data and benchmarked it using injected waveforms. Aurélien J.G. Wyn-gaard wrote subsequent versions of *Lussac*, refined the existing metrics, included the SDR and incorporated the cross-contamination. He implemented the graph analysis approach for the inter-analysis merge and developed the algorithm for identifying and resolving erroneous merges. He created the synthetic dataset and benchmarked all the packages, generating the classification tools used for their comparisons. He wrote this version of the manscript, which was reviewed by all authors. Boris Barbour contributed to the initial conceptualisation of the consensus algorithm and to many decisions of analytical strategy throughout the development of the package.

## Appendix A Synthetic benchmark complete results

Complete results for the analysis of Fig. 4, including all 11 individual analyses, *Lussac* (both heavy and light versions), and a reference are shown in Fig. A1. *Lussac* heavy was the merge of all 11 analyses, whereas *Lussac* light was the merge of 4 pre-defined analyses: Kilosort 2 (optimized), Kilosort 2.5 (optimized), Kilosort 4 (optimized) and SpyKING CIRCUS 2 (default). The reference was created by taking information from the ground-truth data, and selecting the best unit for each neuron in all 11 individual analyses. Note that *Lussac* can outperform the reference, because of the creation of the consensus spike train for each unit, which can improve unit accuracy.

**Supplementary Fig. A1.**
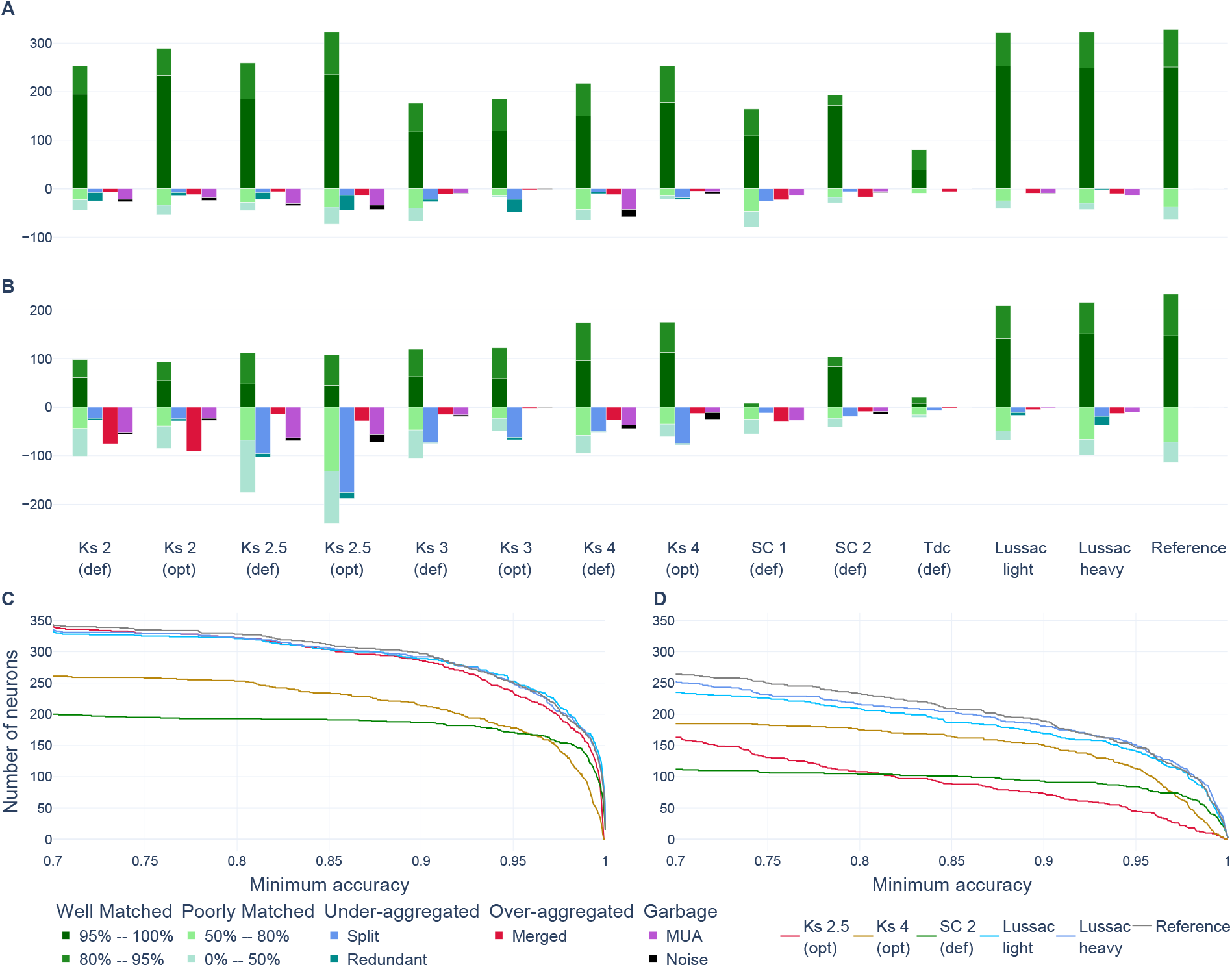
More complete version of the analysis of Fig. 4. (**A-B**) Classification of every unit for every analysis for the stationary (A) and drifting (B) cases. (**C-D**) Number of neurons recovered with an accuracy of at least the value of the *x*-axis for the stationary (C) and drifting (D) datasets. def = default, opt = optimized.

## Appendix B Injection benchmark complete results

Complete results for the analysis of Fig. 6, including all 8 individual analyses and *Lussac* (which is the merge of all 8 analyses) are shown in Fig. B2.

**Supplementary Fig. B2.**
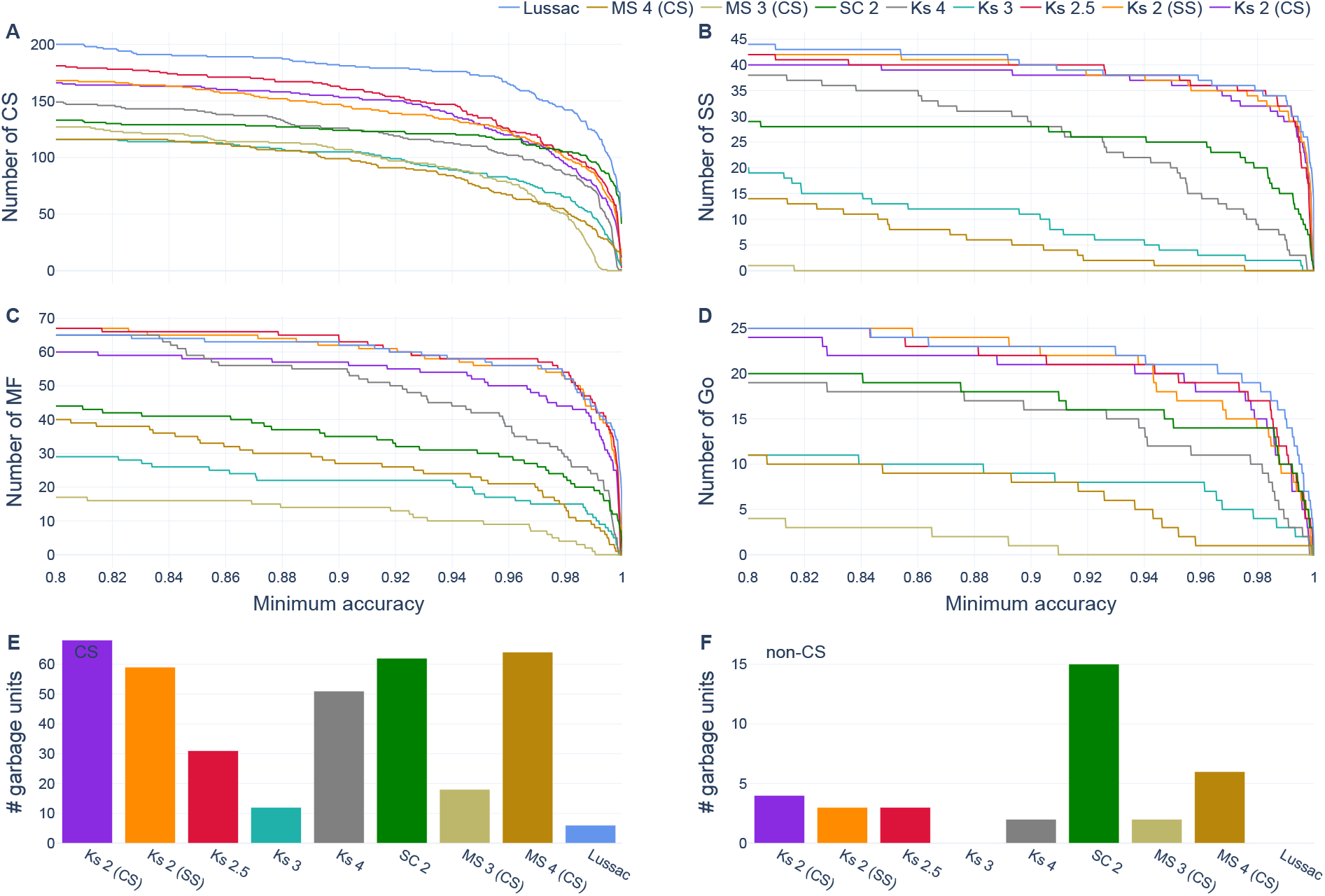
More complete version of the analysis of Fig. 6. Number of injected units recovered with an accuracy greater than or equal to the value on the *x*-axis for complex spikes (**A**), simple spikes (**B**), mossy fibers (**C**) and Golgi cells (**D**). (**E**) Number of garbage complex spike units for each analysis. (**F**) Number of garbage non-complex-spike units for each analysis.

## Appendix C Study of contamination and SD ratio in injected data

For each of the four analyses *Lussac* heavy, Kilosort 2.5 (optimized), SpyKING CIRCUS 2 (default) and MountainSort 3 (SS optimized) and for each dataset, units matching an injected neuron with an accuracy of at least 33 % were retained. For each, the estimated contamination and SD ratio were computed, as well as the real contamination (computed as 1 − precision). It can be seen in Fig. C3 for Kilosort 2.5 and SpyKING CIRCUS 2, that Eq. 2 estimating the contamination is quite accurate, with some variance expected from the randomness of the refractory period violations. However, some units output by MountainSort 3 have significantly lower apparent contaminations than the true values. We believe this may result from the absence of a template-matching procedure to (re)detect of spikes after clustering. This may tend to exclude short-interval spikes whose waveforms interfere with each other from the output clusters. This would deplete spikes specifically from the refractory period and thereby artefactually reduce the apparent contamination. *Lussac* also underestimates the contamination, because, when comparing multiple analyses, it is most likely to output a unit whose contamination is underestimated rather than overestimated.

When comparing the estimated contamination with the SD ratio, is seems that there is an anti-correlation between them. Our hypothesis is that this results from a selection bias: units that have both a high contamination and high SD ratio are easy to detect as bad and never output by the spike-sorting algorithm. This is why the use of both metrics is important.

**Supplementary Fig. C3.**
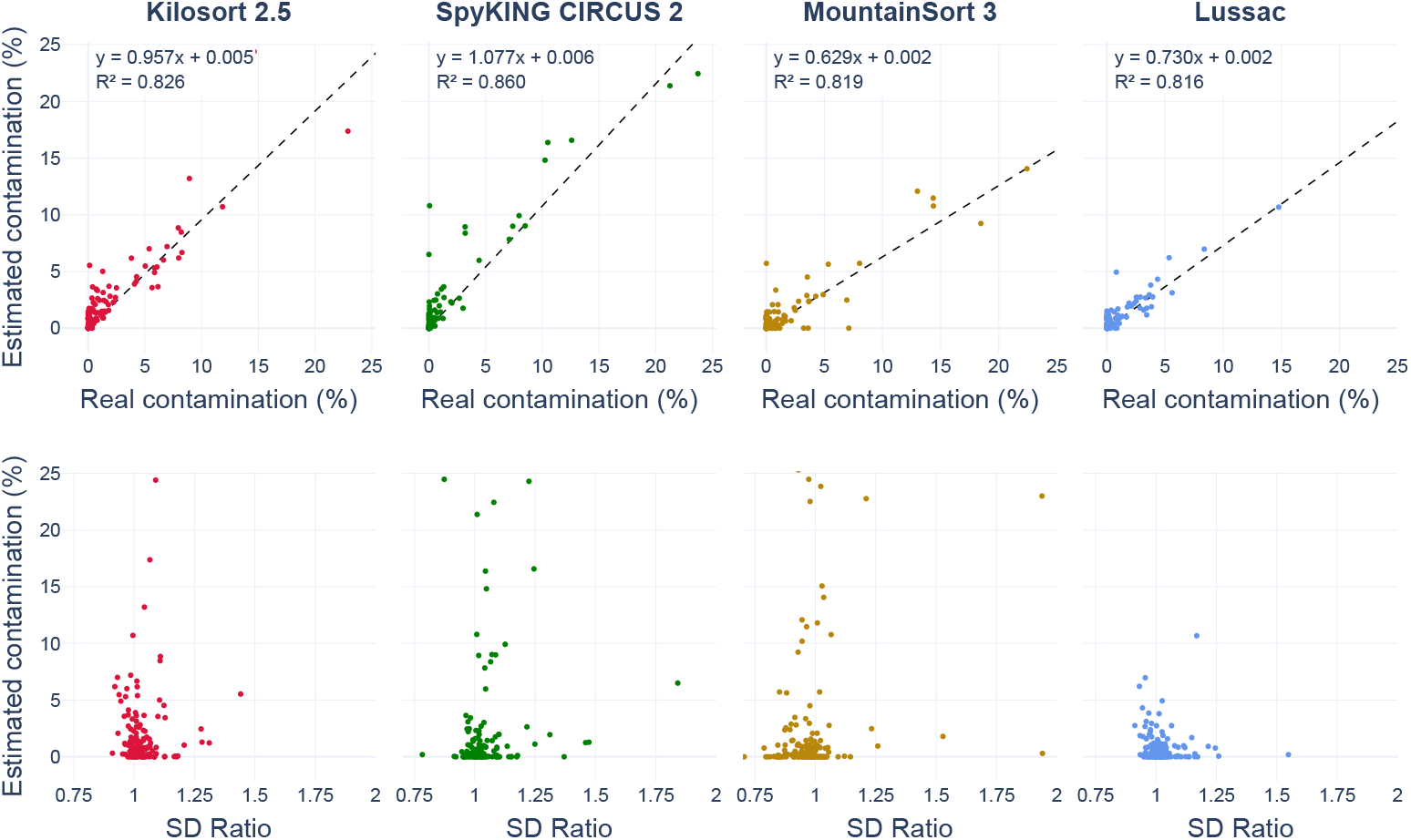
Comparison of the estimated contamination (Eq. 2) against real contamination (top row) and computed SD ratio (bottom row) on the cerebellar injected datasets (all mice).

## Appendix D Study of erroneously merged units

Fig. D4 shows three examples of merged units from the Kilosort 2.5 (default) analysis out of the 6 found in the output for the stationary synthetic dataset. We believe that most experts looking at the available data (without the ground-truth data) would consider these units to be good (expect the first one if one uses the SDR metric). However, the ground-truth data shows that these units are actually merges of 2 (or 3) neurons (that are moreover uncorrelated!). The apparent low contamination is probably due to a poor retrieval of colliding spikes.

Lussac managed to perfectly split cases 1 and 3 (both neurons were outputed with a really high precision (*>*95 %)), while case 2 was partially resolved (neuron #1525 was outputed with high precision, while the others weren’t).

**Supplementary Fig. D4.**
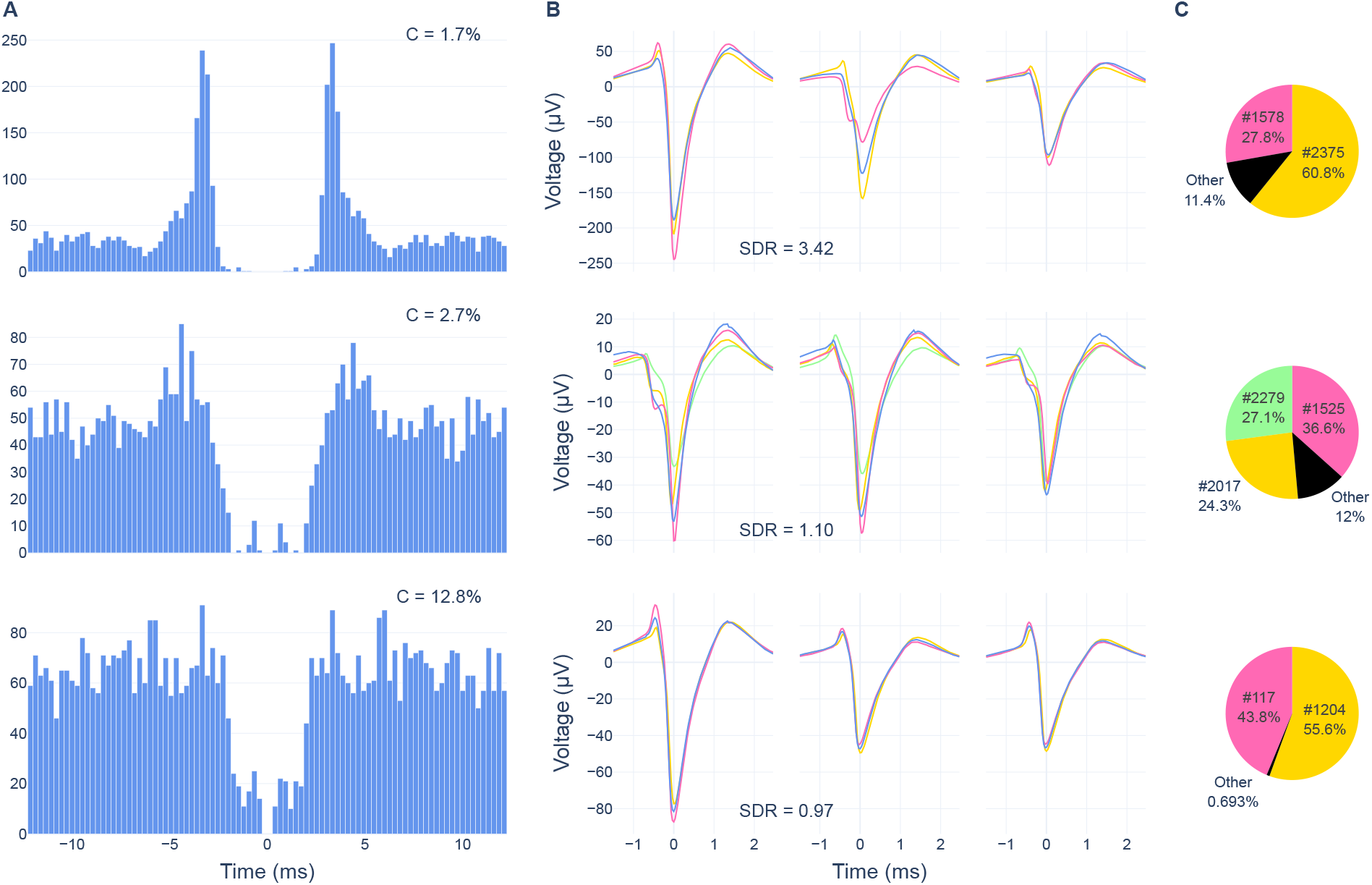
Example of 3 merged units from the Kilosort 2.5 (default) analysis of the stationary synthetic dataset. (**A**) Auto-correlogram of the unit, with its estimated contamination. (**B**) Template of the unit (blue) and of the underlying neurons (other colors) on the 3 best channels. The noted SDR is that of the unit. (**C**) Proportion of spikes in the unit coming from the neurons that were merged and others.

## Appendix E Spatio-temporal profiles of Purkinje cell simple and complex spikes

The extracellular signals of simple and complex spikes of a Purkinje cell are generated through different mechanisms. For simple spikes, the main dipole is generated by the action potential itself in the axon initial segment and soma. Inward membrane current in these compartments generates an extracellular negativity and returns via the proximal dendrites, generating a more diffuse positivity in that region. For complex spikes, the initial driving sink is the large, distributed synaptic current entering the proximal dendrites (where the generated field is called a “fat spike”; [16]), and it appears to return near the soma [17, 18], probably through a somatic potassium conductance. Upon these synaptically-generated dipoles are super-imposed a small number of somatic action potentials, the first of which closely resembles the simple spike, while the remainder appear as “spikelets” that involve either or both of dendritic calcium spikes and axonal action potentials [19, 20].

The waveforms of simple and complex spikes both depend upon the position of the electrode with respect to the Purkinje cell. Examples are shown in Fig. E5A. Although the forms of the complex spike are quite variable (and change sign in some places), the components of the different dipoles will always have similar frequency components, simply because every inward current must always be balanced by an outward current to satisfy conservation of charge. This relative constancy of the frequency components can be seen in the wavelet decompositions (Fig. E5D). The CWTs (Continuous Wavelet Transforms) were produced using the MNE package [21], with the “morlet” wavelet family.

**Supplementary Fig. E5.**
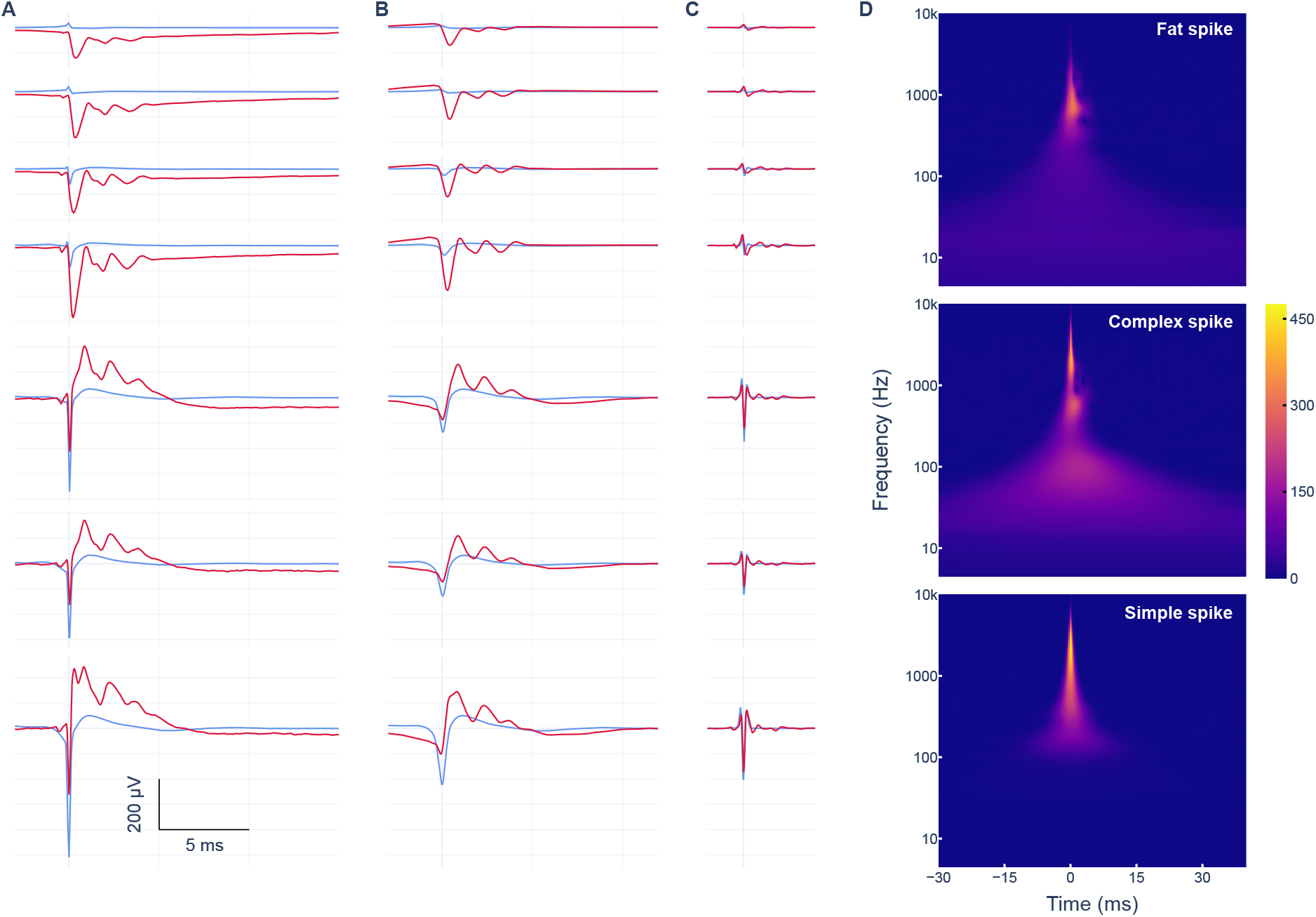
Time-frequency decomposition of simple and complex spike waveforms. Specimen average simple (blue) and complex (red) spike waveforms before filtering (**A**), bandpass-filtered at 60 to 1000 Hz (**B**) or 1.5 to 9 kHz (**C**); recorded on multiple channels (rows). (**D**) Wavelet spectrograms of the dendritic (top, *Fat spike*) and somatic (middle) complex spike waveform, as well as the somatic simple spike (bottom).

The frequency components can also be observed in the temporal domain. Thus, in Fig. E5B,C, we illustrate simple and complex spike waveforms after bandpass filtering with low-(B) and high-(C) frequency ranges. Not accounting for the spikelets, the simple and complex spikes have similar waveforms in the high frequencies (*>*1.5 kHz). This is expected, as these frequencies are largely represented by the somatic action potential, and constitute a verification tool to make sure the simple and complex spike units come from the same Purkinje cell. In contrast, the lower-frequency components are present in the complex spike at a reasonable amplitude at most recording locations.

The nearly ubiquitous low-frequency components exclusive to complex spikes constitute a key characteristic for detecting them and for discriminating them from simple spikes, as has already been implemented [22] and characterized [23]. The lower frequencies are not the only hurdle in correctly detecting complex spikes, as they are also rare (average firing rate around 1–2 Hz), the spikelets render the waveform variable and the presence of a simple spike just before the complex spike can dramatically change the waveform. The challenges are described in depth in [24].

## Appendix F Estimation of noise without spikes from one unit

We demonstrate Eq. 10. Under the assumption of independent random variables for the template *W* and the noise *N*, then their variances sum. However, the template isn’t always continuously present, because spikes only arise intermittently. Indeed, for a unit of template width *w* with *N* spikes, the templates span a fraction 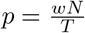 of the trace, where *T* is the duration of the recording. Hence, we can create a template trace *W*^′^ containing zeros everywhere and the *N* non-overlapping templates. In this case, the variance is:

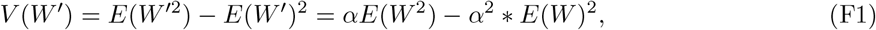

where *E* is expectation, *V* variance and *α* the fraction of the recording spanned by the waveform templates. We subtract this result from the variance of the whole trace to obtain the variance of the underlying noise (Eq. 10)

## Appendix G Contamination

Here we prove Eqs. 1 and 2. The goal is to estimate the contamination 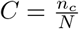 of a given spike train with *N* = *n*_*t*_ + *n*_*c*_ true and contaminant spikes, under the assumption of stationary firing statistics.

Under the model of contaminant spikes coming from a single neuron (*C*_1_) with its own refractory period of at least *t*_*r*_, refractory period violations can only occur between a true and a contaminant spike. Under the hypothesis that both neurons are uncorrelated, the number of violations closely follows a Bionomial distribution, where each (true, contaminant) pair of spikes has a probability 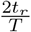 of being separated by *t*_*r*_ or less (ignoring border effects). There are *n*_*t*_ × *n*_*c*_ such pairs, giving an expected total number of violations of:

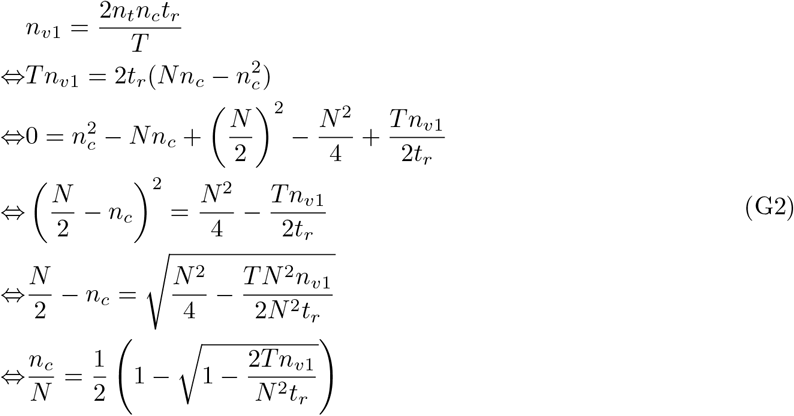

where the the negative root has been eliminated in the penultimate line because 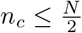.

The model of completely random contaminant spikes (*C*_∞_) is very similar, the difference being that refractory period violations between contaminant spikes are also possible. Since the spikes occur at random, the number of violations follows a Binomial distribution, with probability 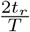 and number of pairs 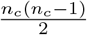. Hence, the expected total number of violations is:

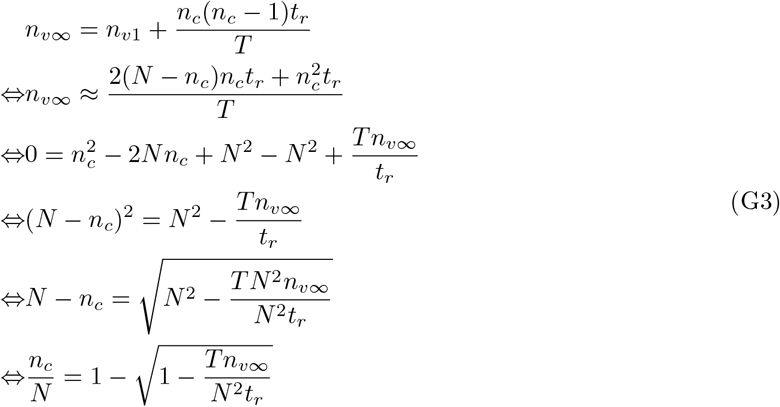

where the the negative root has been eliminated in the penultimate line because *n*_*c*_ *< N*.

## Appendix H Invariance of correlogram expectation under random sampling

Both auto- and cross-orrelograms are powerful diagnostic tools for post-processing in spike-sorting. One key feature of correlograms that we prove next is the fact that the correlogram of a random sample of spikes from the original train will have the same expected shape as the original correlogram.

For a given spike train of a parent unit *A*, its auto-correlogram is represented by *C*_*A*_. We *randomly* sample the spikes of *A* without replacement, such that each spike independently has a probability *p* of being selected to create the child unit *B*. As the process is random, we cannot know exactly the auto-correlogram of *B*, but we can compute its expectated value *C*_*B*_.

For each bin *δt*, there are *C*_*A*_(*δt*) pairs of spikes. However, these pairs are **not** independent, and neither are the spikes (because a spike could contribute to two pairs, or even more). There are thus *n* spikes, each contributing to *n*_*k*_ pairs. If we add the pairs from each spikes, then we get twice the total number of pairs: 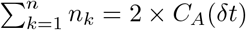

After sub-sampling, each spike independently has a probability *p* of being retained. If it is, then every other spike has, independently, a probability *p* of also being retained. For this initial spike, the number of pairs after sampling thus follows a binomial distribution, whose mean is *n*_*k*_ × *p*. The new number of pairs becomes:

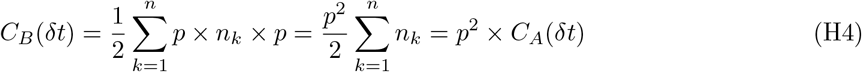

Thus, the auto-correlogram *C*_*B*_ is a scaled version of *C*_*A*_, with a scale factor of 1*/p*^2^. (If the correlogram is calculated at zero shift, at this shift only a different scaling of 1*/p* applies.)

We can also compute the cross-correlogram between units *B* and *C*, resulting from two random samples from unit *A*. Depending on the sampling method, this could represent a split, or partially overlapping units. We can follow the same reasoning as previously, except that the probability of a spike being in *C* is *q* rather than *p*. We thus have the formula for the cross-correlogram:

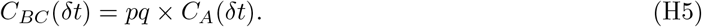

## Footnotes

1 **L**ots of **u**nique **s**pike-**s**orting **a**nalyses **c**ombined

2 A rigid drift is a drift that is the same along the whole probe, which isn’t necessarily the case as the brain is a soft tissue [1]

